# Poly-Target Selection Identifies Broad-Spectrum RNA Aptamers

**DOI:** 10.1101/302745

**Authors:** Khalid K. Alam, Jonathan L. Chang, Margaret J. Lange, Phuong D.M. Nguyen, Andrew W. Sawyer, Donald H. Burke

## Abstract

Aptamer selections often yield distinct subpopulations, each with unique phenotypes that can be leveraged for specialized applications. RNA aptamers that bind HIV-1 reverse transcriptase (RT) exhibit potent RT inhibition and suppress viral replication when targeting the strain-specific RT that they were originally selected to bind, but some of these same aptamers fail against single-point mutant and phylogenetically-diverse RTs. We hypothesized that a subset of the total aptamer population in libraries pre-enriched against a single RT may exhibit broad-spectrum RT binding and inhibition, and we devised a multiplexed Poly-Target selection approach to elicit those phenotypes against a panel of diverse primate lentiviral RTs. High-throughput sequencing of starting, negative, and final libraries, followed by analysis of coenrichment and codepletion in parallel and duplicate selection trajectories, narrowed the list of candidate aptamers by orders of magnitude. Biochemical characterization of candidates identified a novel aptamer motif and several rare and unobserved variants of previously-known motifs that inhibited recombinant RTs from HIV-1, HIV-2 and SIV to varying degrees. These broad-spectrum aptamers also suppressed replication of viral constructs carrying phylogenetically-diverse RTs. The Poly-Target selection and coenrichment approach described herein is a generalizable strategy for identifying broad-spectrum behavior and cross-reactivity among related targets from combinatorial libraries.

## INTRODUCTION

Aptamers are single-stranded nucleic acids that bind with high affinity to defined molecular targets, and they can be generated through an iterative *in vitro* selection process known as SELEX (1–3). While the aptamer field has traditionally sought to maximize selectivity and molecular discrimination among related molecular targets, aptamers with broad target recognition can be especially useful for recognizing entire classes of chemically-related metabolites or evolutionarily-related protein families. In the present work, we explore this theme by seeking aptamers that recognize increasingly divergent representatives of the primate lentiviral family of reverse transcriptases.

RNA aptamers selected against HIV-1 reverse transcriptase (RT) are able to bind RT and inhibit its enzymatic activities, and they overwhelmingly form pseudoknot structures (4–7). These pseudoknot RNA aptamers show robust *in vitro* inhibition of the RT they were selected to bind (8) and they suppress viral replication and infectivity in cell culture (9–13). Structural and biophysical investigations suggest that these pseudoknot RNA aptamers bind the closed conformation of RT in a positively-charged groove, 50-60 Å in length, that spans the polymerase and RNase H domains (14, 15). This same groove can accommodate 16-19 base pairs of nucleic acid and naturally binds a duplex of viral RNA and host tRNA_Lys_ primer. An observation from our lab is that a common polymorphism in this groove, near the polymerase active site (K277 in the p66 subunit), confers resistance to the Family 1 Pseudoknot (F1Pk) subset of RNA aptamers (16), thereby preventing broad-spectrum inhibition. In contrast, singlestranded DNA aptamers selected against RT (17) demonstrate broad-spectrum inhibition *in vitro* (18–21), but their use in cell culture assays of viral replication or *in vivo* is complicated by the requirement to produce or deliver single stranded DNA in the relevant cells.

High-throughput sequencing of anti-RT RNA aptamer libraries recently revealed several additional motifs capable of RT binding and inhibition, including the (6/5) asymmetric loop ((6/5)AL) (7) and the UCAA bulge (22). These non-pseudoknot motifs were difficult to detect with low-throughput sequencing, as their low sampling frequencies were eclipsed by the abundance of aptamers that carried F1Pk or F2Pk motifs (>93% in one library (7)). Although these novel motifs were neither selected nor initially characterized specifically for their broad-spectrum inhibition, recent work, done in conjunction with the work described here, demonstrated that these non-pseudoknot motifs were capable of broad-spectrum suppression of viral replication when actively encapsidated into nascent virions (13). These results indicate that even within a highly evolved library, there remains an important untapped phenotypic diversity that is difficult to capture by conventional selection strategies. The central goal of the present work is to demonstrate a general method that can be used to find aptamer subsets with broad-spectrum target recognition for any family of proteins. The additional functional diversity present in the RT-binding libraries suggested that new genotypes and phenotypes were potentially accessible under different selection conditions and made them ideal for demonstrating the approach.

Herein we describe a Poly-Target selection method to enrich for RNA aptamer subpopulations with broad-spectrum binding and inhibition of diverse primate lentiviral RTs. Rather than performing a *de novo* selection from a random-sequence library, we utilized two well-characterized libraries that had been pre-enriched for binding RT from a specific HIV-1 strain. Negative selections against nitrocellulose were performed to remove RT-independent binders, followed by three rounds of positive selection against a panel of seven diverse primate lentiviral RTs, in parallel. Each selection trajectory was performed in duplicate, and high-throughput sequencing of the starting, negative, and final libraries enabled an informatics approach to rapidly identify candidate aptamers through coenrichment analysis of the various selection trajectories. Structural and biochemical characterization of candidates identified several rare variants of known motifs and one previously unknown motif capable of varying degrees of broad-spectrum inhibition of RT polymerase activity. Furthermore, these broad-spectrum aptamers inhibited RTs from Subtype C and other clades that were not included in the selection panel and suppressed viral replication of constructs containing diverse RTs. We believe the Poly-Target selection method is broadly applicable to any aptamer or combinatorial selection approach that seeks to identify cross-reactivity.

## RESULTS

### Poly-Target Selection Panel

Two distinct, pre-enriched aptamer libraries with randomized regions of either 70 or 80 nucleotides (70N and 80N, respectively) were used to initiate Poly-Target selection. These libraries were originally selected *in vitro* against recombinant RT from a group M, subtype B strain of HIV-1 (BH10) (6). In these original selections, partitioning for protein-bound RNA was accomplished through eleven rounds of nitrocellulose filtration and three rounds of native gel shifts, for a total of fourteen rounds. The vast majority of RNA aptamers in these populations form pseudoknot structures, with much lower frequencies of other, previously undetected motifs, such as the 6/5 asymmetric loop ((6/5)AL) (7) and UCAA bulge (22). All of these motifs bind with low nanomolar affinity and inhibit RT by competing with primer-template for access to the enzyme. The presence of additional, low-abundance, high-affinity aptamers suggested that significant diversity exists within these Round 14 libraries. A central goal of the present work was to exploit untapped diversity within these pre-enriched libraries by identifying low-abundance structural motifs that may offer broad-spectrum binding and inhibition of RTs.

To elicit broad-spectrum phenotypes from the pre-enriched libraries, we assembled a panel of phylogenetically diverse primate lentiviral RTs (16) (**Supplementary Figure 1**). The panel of targets included RTs from multiple strains of the major group (group M) of HIV-1, which is responsible for the global pandemic and is further divided into subtypes and circulating recombinant forms (CRFs) that contain genomic segments from two or more subtypes. HXB2 is a group M, subtype B strain closely related to the original selection target (98.9% by amino acid identity), and it is often used as a reference strain for sequence comparisons. We therefore included RT from strain HXB2 in the panel as a “continuity control.” Also included was an R277K single point mutant of HXB2, a natural polymorphism which confers resistance to inhibition by F1Pk aptamers *in vitro* (16) and in cell culture (13), to enrich for aptamers that bind in spite of this escape mutation (henceforth referred to as R277K). RT from strain 94CY was included as a representative of subtype A HIV-1, along with RT from 93TH, which groups phylogenetically within CRF01_AE, a circulating recombinant form that carries genomic segments corresponding to subtypes A and E.

The selection panel also included RTs from HIV-1 strains from outside Group M and from simian immunodeficiency virus (SIV_cpz_Pts). Although these additional strains are of less epidemiological relevance, they help to capture the broader genetic diversity of primate lentiviruses (**Supplementary Figure 1**). The outlier group of HIV-1 (group O) is phylogenetically distinct from group M and was therefore represented in the selection panel by an RT from the group O strain MVP. Because HIV-1 arose from zoonotic transmission of SIV_cpz_ from chimpanzees (23), we included an RT from the TAN1B strain of SIV_cpz_. Finally, we included RT from the EHO strain of HIV-2, which is only distantly related to HIV-1, to further broaden the scope of the Poly-Target selection. Selecting independently against each of these RTs presents a gradient of selection pressures on the pre-enriched library to search sequence space for ligands that are capable of making molecular contacts with conserved features of primate lentiviral RTs. A table depicting the unabbreviated names of the strains, their reference sequences, and amino acid changes relative to the database reference sequences are provided in **Supplementary Table S1**.

### Poly-Target Selection, High-throughput Sequencing and Bioinformatics

A schematic for the Poly-Target selection strategy is shown in **Figure 1**. Pre-enriched Round 14 libraries (70N and 80N) were transcribed and partially depleted for nitrocellulose-binding species. Libraries were then separated into replication populations (A and B) and further split into independent populations for each of the seven RT targets. For each round of Poly-Target selections, transcribed RNA for a given trajectory was independently incubated with a single RT from the selection panel. Bound RNA:RT complexes were captured on nitrocellulose filters, and the recovered RNA was reverse transcribed into cDNA and then PCR amplified for additional rounds of selection or for high-throughput sequencing (HTS). Each selection trajectory was performed for a total of three rounds, after which the starting Round 14 libraries, the nitrocellulose binding populations, and the Round 17 libraries were submitted for HTS.

**Figure 1.**
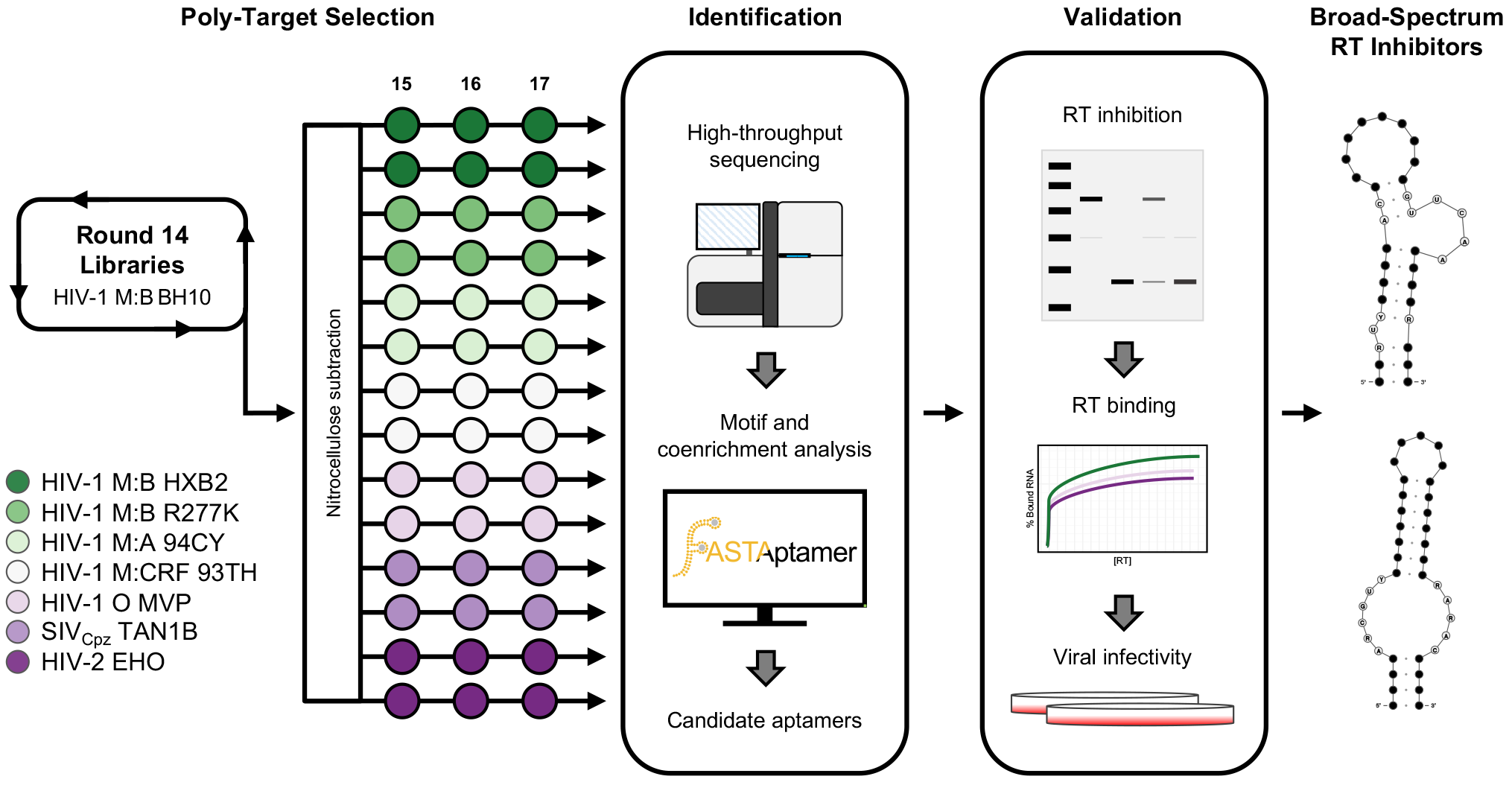
A Poly-Target selection strategy to identify broad-spectrum RT inhibitors. In the Poly-Target selection approach a pre-enriched aptamer library is subjected to additional rounds of systematic evolution of ligands by exponential enrichment (SELEX) against a panel of related targets along separate trajectories. In this work, 70N and 80N libraries that had been pre-enriched through 14 rounds of binding to an RT from HIV-1 M:B (BH10) (13) were first subjected to a negative selection (nitrocellulose subtraction) prior to three additional rounds of independent selections, in duplicate, against a panel of phylogenetically-distinct RTs from various strains of HIV-1, HIV-2 and SIV. Following the Poly-Target selection, high-throughput sequencing and coenrichment analysis were used to identify candidate broad-spectrum aptamers for downstream biochemical and biological validation.

HTS of the libraries generated over 184 million raw reads. Demultiplexing, trimming and quality filtering (**Supplementary Figure S2**) of sequences left over 52 million high quality reads for the 70N library and 54 million for the 80N library. Each selection trajectory contained an average of three million total reads and 35,000 to >100,000 unique sequences (**Supplementary Table S2**). The average number of reads per unique sequence increased for all populations, indicating enrichment and convergence on sequence space within each library, with the exception of 70N TAN1B (A), which was the only population with fewer than one million total reads (**Supplementary Figure S3**). Aptamer populations were further processed by the FASTAptamer toolkit to determine sequence frequencies, compare populations and calculate enrichment of each unique sequence relative to the starting Round 14 library (24). Sequences were clustered with FASTAptamer using Levenshtein edit distance - a string similarity metric that calculates the minimum number of insertions, deletions and substitutions necessary to transform one sequence into another. Sequences within a small edit distance from each other likely diverged from a parent sequence in an earlier population and grouping them into a single cluster simplifies analysis. Based on previous observations that sequences in these populations typically diverge by less than 10% from their parent sequences (7), we defined clusters as having a maximum edit distance of 7 for the 70N library and 8 for the 80N library.

To determine the degree to which replicate selections against the same target experienced similar evolutionary paths, normalized sequence frequencies from the A and B trajectories for each target were compared using FASTAptamer-Compare (24). For both the 70N and 80N Round 14 libraries, we observed a tight *y* = *x* relationship between replicates (**Supplementary Figures S4A and S5A**), suggesting highly similar input diversity for each pair of replicate trajectories for a given target. In contrast, comparing sequence frequencies between replicate trajectories after three rounds showed a multimodal distribution for many populations, wherein different subsets of sequences either enrich, remain neutral, or deplete (**Supplementary Figures S4B-H and S5B-H**). These minor variations between replicate trajectories likely reflect stochastic variations in sample handling (25) and highlight the potential value in providing multiple, independent evolutionary opportunities for a given sequence to enrich or deplete. As expected, distributions were widened more dramatically when comparing different selection rounds or selections against different targets (**Supplementary Figures S6 and S7**).

### Coenrichment Analysis Identifies Candidate Broad-Spectrum Aptamers

Sequences from the Poly-Target selection that enriched in more than one selection trajectory were expected to be the most promising candidates as broad-spectrum aptamers. As a first step towards coenrichment analysis, we identified sequences within each post-selection population that met a given enrichment threshold, which we defined as a two-fold or greater increase in reads per million (RPM) relative to the starting Round 14 population (**Supplementary Figure S8A**). To mitigate the potential impact of low abundance sampling artifacts, we confined this analysis to sequences with an aggregate RPM ≥ 10 when summed across the populations being compared (e.g., Round 14 RPM + Round 17 RPM). We also included sequences that were below the detection limit in Round 14 libraries but later increased in relative abundance and were sampled with ≥ 10 RPM in Round 17, although their absence in Round 14 precluded calculating an enrichment ratio. On average, each trajectory had approximately 1,800 sequences that met these enrichment criteria (**Supplementary Table S3**). Depleting sequences were similarly flagged by identifying sequences with aggregate RPMs of ≥ 10 that either decreased by at least two-fold (0.5-fold enrichment) in each trajectory, or that were present in Round 14 at ≥ 10 RPM and below the detection limit in Round 17. Each trajectory had approximately 2,600 sequences that met these depletion criteria (**Supplementary Table S3**). In nearly all trajectories, the number of sequences that depleted throughout the course of the selections exceeded the number that enriched.

We reasoned that broad-spectrum aptamers should enrich along multiple selection trajectories, while specialist aptamers should deplete in most or all trajectories. Therefore, we next identified the sets of sequences that experienced coenrichment or codepletion in two or more trajectories (**Supplementary Figure S8B**). These were pooled according to starting library and replicate, irrespective of which individual trajectories provided the basis for identifying them as coenriching or codepleting. Each starting library (70N and 80N) and each replicate within them (A and B) were analyzed independently, thereby generating four unique sets of coenriched sequences (average of 3,200 per set) and four sets of codepleted sequences (average of 3,500 per set) (**Supplementary Table S4**). Sequences demonstrating coenrichment or codepletion in the negative nitrocellulose selections were eliminated from analysis, reducing each library replicate to an average of 2,500 coenriched and 1,800 codepleted sequences. Dataset complexity was further reduced by reclustering each set of coenriched or codepleted sequences into sequence families. To facilitate comparison across trajectories, the identified sequences were mapped to their cluster identity from Round 14 (Supplementary Figure S9). Clustering and mapping in this manner allowed us to readily discern highly-enriched clusters that arose in either or both replicate trajectories.

As individual clusters can contain numerous sequence variants, individual sequences within those clusters can sample advantageous or deleterious mutations. To focus on clusters where the majority of sequences exhibited coenrichment with minimal codepletion, we coupled the output from the coenrichment and codepletion analysis (Supplementary Figure S10). Fourteen clusters from the 70N libraries and nineteen clusters from the 80N libraries were chosen for further characterization based on this analysis (33 total clusters). Two clusters that heavily depleted in the 70N trajectory (70N 2 and 70N 3) were also included to test whether codepletion is predictive of low fitness. For each cluster, the most abundant corresponding sequence in Round 14 was chosen to represent the entire cluster in biochemical characterization. Candidate aptamers are named by their library of origin and their cluster identity in the pre-enriched Round 14 libraries (*e.g.* aptamer 70N 89 is the 89^th^ most abundant duster defining sequence in the 70N Round 14 library). Sequences that were not sampled in the Round 14 libraries were named according to the cluster identity generated in the coenrichment analysis. A heat map of genotypic frequencies for each selected candidate aptamer in each trajectory is shown in **Figure 2A**, with more detailed information on the frequencies of individual sequences in each trajectory detailed in **Supplementary Table S5**.

**Figure 2.**
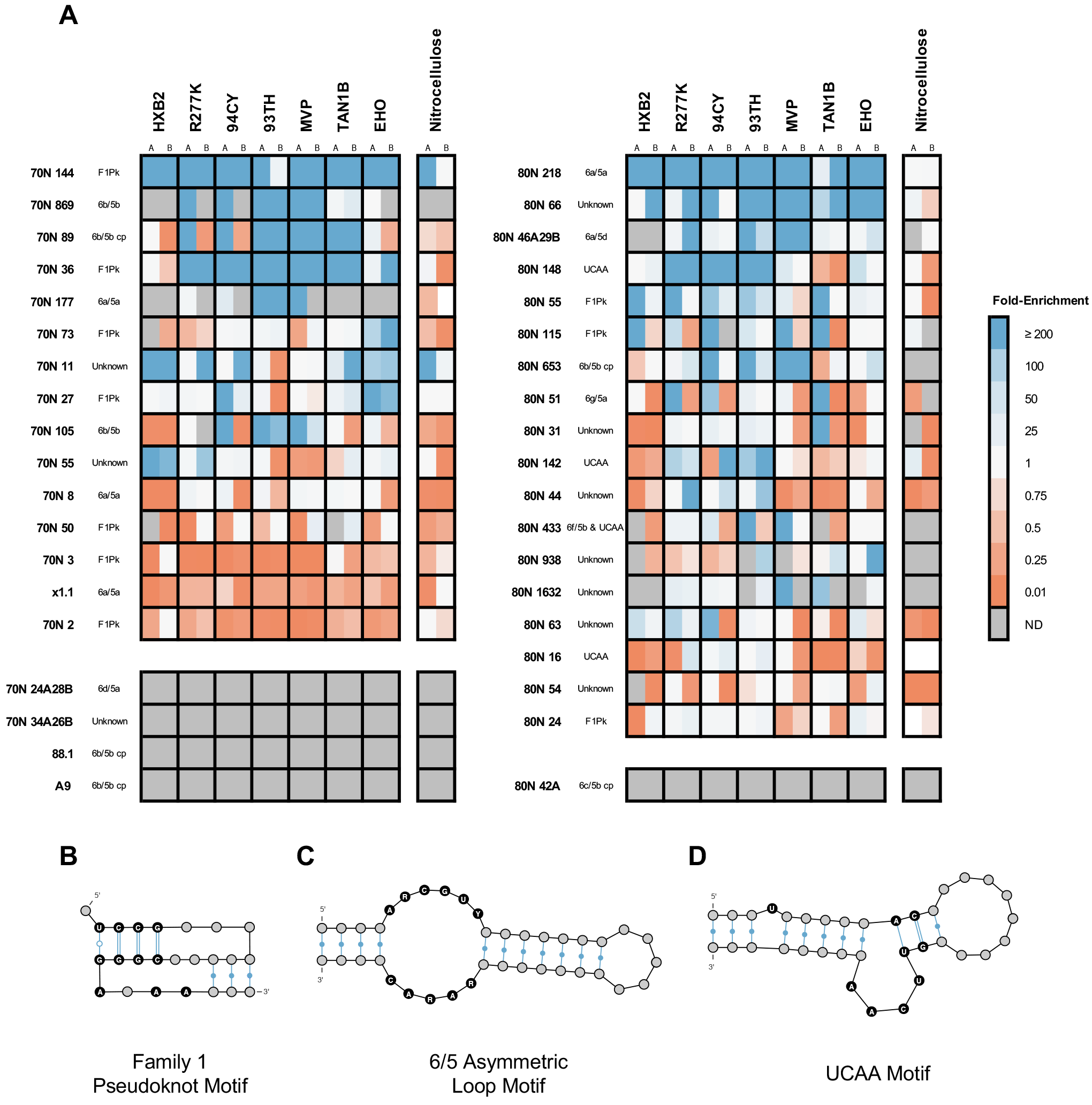
Coenrichment analysis identifies candidate broad-spectrum aptamers. (**A**) A heatmap of sequence fold-enrichment from the round 14 starting library of candidate aptamers depicts enrichment (blue) or depletion (orange) in each replicate selection trajectory (A or B) and in the nitrocellulose binding population. Aptamers are grouped by 70N (left set) or 80N (right set) and are sorted top-to-bottom by decreasing enrichment, averaged across all RT targets. Sequences included in the characterization but that were not detected (ND) in either round 14 or 17 and therefore did not have a enrichment value are colored in grey. Motif names are included to the right of the aptamer identity. Schematic depictions of the secondary structures of the major RT-inhibiting motifs: (**B**) Family 1 Pseudoknot (F1Pk), (**C**) 6/5 Asymmetric Loop ((6/5)AL) and (**D**) UCAA. Secondary structures generated using VARNA (58).

### Genotype Analysis Identifies Known, Rare and Previously Unobserved Sequence Motifs

The 33 sequences identified above (**Figure 2, Supplementary Figure S9**, Supplementary Table S5) and two additional sequences that both heavily codepleted were analyzed for the presence of sequence motifs that have been previously shown to robustly inhibit subtype B RTs *in vitro* and to suppress viral replication in cell culture (7, 9–13, 22). The FASTAptamer-Search function of the FASTAptamer toolkit (24) was employed to search for degenerate sequence motifs, and Mfold (26) was used to generate secondary structure predictions. Although Mfold is unable to predict pseudoknot motifs, we reasoned that it would help to provide structural evidence for UCAA and (6/5)AL motifs and could help identify aptamers with novel motifs. Applying this approach with relaxed search constraints identified the three major secondary structural motifs known to bind HIV-1 RT – pseudoknot, (6/5)AL and UCAA – within most, but not all of these 35 sequences.

FASTAptamer-Search identified ten sequences as F1Pk (**Supplementary Table S6**), including the two 70N sequences noted above as having codepleted significantly (70N 2 and 70N 3). The F1Pk motif is characterized by the string UCCG-N_7/8_-CGGGANAAN_≥3_, where N refers to any nucleotide, and N_≥3_ refers to a segment that base pairs with at least three connector nucleotides to form Stem II (**Figure 2B**) (4–7). Some of these F1Pk-like aptamers diverged slightly from the expected sequence motif definition, and their enrichment patterns appear to reflect their specialized characteristics. For example, 70N 27 and 70N 73 contained only two potential base pairs in Stem II. They also showed the strongest enrichment in the EHO (HIV-2) trajectory, potentially allowing for this variation of the F1Pk motif to maximize contacts with that particular RT. Similarly, while nine of the ten F1Pk sequences contain a seven or eight nucleotide connector, 70N 144 contains a ten-nucleotide connector. The strong enrichment of this sequence in all of the trajectories, including the nitrocellulose binding population, indicates that it may be a selection parasite, rather than a high-affinity RT binder.

The UCAA motif is minimally defined by a four-nucleotide UCAA bulge flanked by AC/GU closing base pairs on one end and by a stem that is interrupted by a single unpaired U on the other end (**Figure 2D**) (22). Searching for the UCAA motif identified four sequences, two of which (80N 16 and 80N 148) fully conformed to the motif definition. 80N 16 is identical to aptamer 80.103, which had been identified previously and shown to inhibit HXB2 RT *in vitro* and to suppress viral replication in cell culture (22). The core element within 80N 148 was validated through aptamer truncation and RT inhibition analysis (**Supplementary Figure S11**). The UCAA elements in the other two identified aptamers were within non-canonical structural contexts and may only superficially resemble the UCAA motif. Aptamer 80N 142 contained the sequence UCAACU but presents the internal CAAC as the four-nucleotide bulge, rather than the UCAA portion, and both flanking U’s are predicted to pair within stems. Nevertheless, this sequence also contained the downstream unpaired U and upstream AC/GU base pairs characteristic of the motif. Although the fourth sequence (80N 433) contained a UCAA motif, it was predicted to present this sequence either as a tetraloop capping an interrupted helix, or as a part of a seven-nucleotide bulge (GUUCAAA), neither of which conforms to the canonical structural definition of the UCAA.

The (6/5)AL motif is characterized by the presence of six unpaired nucleotides (ARCGUY) opposite five unpaired nucleotides (RARAC) in an asymmetric loop embedded within a stem (**Figure 2C**); where R is any purine and Y is any pyrimidine. Of the 16 loop combinations that fit this definition, 6a/5a (AACGUU/GAAAC) and 6b/5b (AGCGUC/AAGAC) variants accounted for >98% of all (6/5)AL motifs in the 70N Round 14 library, along with a small fraction of additional minor variants (7). Searching the candidate sequences for the (6/5)AL motif identified eight sequences that fit the consensus. The 6a/5a variation was present in three sequences (70N 8, 70N 177, and 80N 218). Although most of the previously-identified (6/5)AL aptamers utilize portions of the 80N constant region to form the motif (7), 80N 218 contained the motif entirely within the aptamer’s randomized region. Four aptamers containing the 6b/5b variation of the motif were also identified, with two in the common circularly permuted form (70N 89 and 80N 653) and two in the less-common, non-permuted form (70N 105 and 70N 869). A circularly permuted 6c/5b variant that contained the motif within the randomized region was also found (80N 42A). The 6c/5b combination is exceedingly rare, having been previously observed at less than 0.1% of the 70N Round 14 population (7). To validate these motif identifications, RNA 3’ boundary determination for the aptamer:RT complex and inhibition assays with truncated variants were carried out for two non-circularly permuted (70N 105 and 70N 869) and one circularly-permuted (70N 89) 6b/5b aptamer. All three fully inhibited rT from HIV-1 strain HXB2 when the 3’ end was truncated to only three base pairs beyond the asymmetric loop, irrespective of whether the closing stem was Stem I or Stem II (**Supplementary Figures S12–S14**).

Sequences with no obvious motifs were analyzed for their predicted secondary structures. Four of these contained asymmetric internal loops suggestive of (6/5)AL variants that did not conform to the ARCGUY/RARAC consensus sequence and hence escaped detection in the initial sequence-based screening. Two of these contain known, but extremely rare, variations of the motif, including 6a/5d (AACGUU/<u>U</u>AAAC, non-conforming deviations underlined) in 80N 46A29B and 6d/5a (AACGU<u>G</u>/GAAAC) in 70N 24A28B. These sequence combinations were previously detected in a 70N library with frequencies of less than 0.6% (6a/5d) and 0.08% (6d/5a) of all (6/5)AL reads (7). The other two contained previously unobserved variants of the (6/5)AL motif, including the newly-termed 6f/5b (AGCGU<u>A</u>/AAGAC) in 80N 433 – which is flanked by AG/CC closing pairs in Stem I – and 6g/5a (AGC<u>C</u>UC/GAAAC) in 80N 51. The observation of a variant (6/5)AL motif in 80N 433 was especially surprising, as this sequence also contains a non-canonical UCAA bulge, as described above. Minimization of the 80N 433 aptamer, followed by deletion of either the (6/5)AL motif or UCAA bulge, suggested that only the (6/5)AL motif is required for inhibition of HXB2 RT and that the portion containing UCAA is not sufficient for inhibition (**Supplementary Figure S15**).

In total, motif analysis of coenriched and codepleted cluster seed sequences, combined with biochemical validation of a subset of these, identified ten putative F1Pk aptamers, three potential UCAA motif aptamers and twelve (6/5)AL aptamers that include several circular permutations and rare or novel variants. The F1Pk motif comprised the vast majority of the Round 14 libraries and continued to be a major component of the final populations despite their overall depletion. One aptamer (80N 433) contained both a cryptic (6/5)AL motif and a dispensable, non-canonical UCAA bulge. Ten additional aptamers neither conformed to previously-defined motifs nor showed sequence or structure similarity with each other. Because many of the motif-unknown sequences were also abundant in the nitrocellulose binding populations, some of these aptamers may be low-fitness species with protein independent enrichment. Overall, Poly-Target selection and coenrichment analysis helped to reduce the lead candidate space by several orders of magnitude to identify highly coenriched, rare, and unique genotypes that were present within the pre-enriched Round 14 libraries. It is unlikely that low-throughput sequencing would have identified these unknown and previously unobserved sequences, as their relative abundance in each library remained low even after three additional rounds of selection.

### Broad-spectrum RT Inhibition Varies with Aptamer Motif

To test whether coenrichment against multiple, divergently-related protein targets was an indication of broad-spectrum recognition across the panel of RTs, we assayed the 35 cluster-representing aptamers for inhibition of the DNA-dependent DNA polymerase (DDDP) activity of RT using primer extension assays. Three previously characterized 70N (6/5)AL aptamers (88.1, A9, ×1.1) were included as inhibition controls. Aptamers A9 and 88.1 contain a circularly permuted 6b/5b motif and enriched along many selection trajectories, despite being notably absent from sequences obtained here for the Round 14 library. Aptamer ×1.1 contains a 6a/5a motif and was the most abundant sequence in the 70N Round 14 library (7), yet it was among the most depleted sequences in the Round 17 populations. The vast majority of aptamers containing known motifs robustly inhibited RT from strain HXB2 (**Figure 3A**, top), which is very closely related (98.9%) to the BH10 strain used to pre-enrich these libraries originally (6). In contrast, only three of the aptamers without recognizable structural motifs inhibited RT from HXB2 (70N 11, 70N 55 and 80N 44), suggesting that the other candidates with unknown structures may have been enriched for binding to nitrocellulose, that they bind RT surfaces that do not interfere with DNA polymerization, or that they are selection parasites.

**Figure 3.**
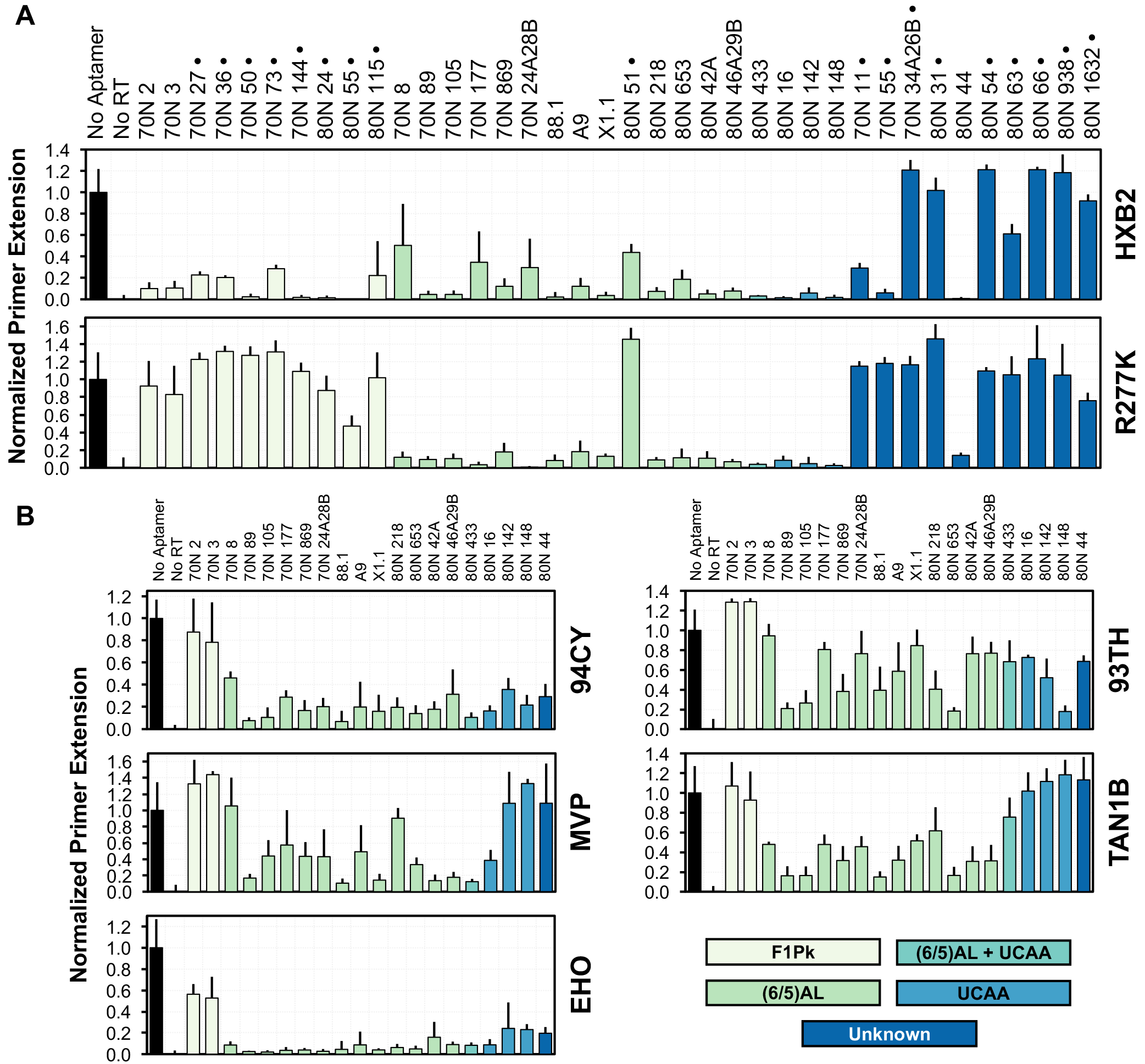
RT inhibition assays reveal broad-spectrum phenotypes. (**A**) Aptamer mediated inhibition of the DNA-dependent DNA polymerase activity of RT was determined through primer extension assays against HXB2 RT (top) and an R277K point mutant of HXB2 RT (bottom) for candidate aptamers identified through coenrichment analysis. Aptamers that failed to inhibit both RTs after two independent measurements (marked **·**) were eliminated from further examination, with the exception of 70N 2 and 70N 3. (**B**) Primer extension assays against non-Group M:B RTs in the selection panel reveal varying broad-spectrum inhibition. Columns are colored according to aptamer motif. Error bars indicate range for samples assayed in duplicate (marked **·**), whereas all other samples are n ≥ 5 and error bars indicate standard deviation.

The R277K point mutant of RT from HXB2 served as initial surrogate for evaluating broad-spectrum inhibition. The K277 polymorphism is prevalent among sequenced strains and was present in five of the seven RTs used in these selections, so we reasoned that true broad-spectrum aptamers should be able to inhibit the R277K RT. When all 38 aptamers were tested for inhibition of R277K, 18 of these inhibited both HXB2 and R277K (**Figure 3A**, bottom). The (6/5)AL and UCAA motif aptamers demonstrated strong inhibition of R277K, with the exception of 80N 51, which weakly inhibited RT from HXB2 and failed to inhibit the point mutant. This aptamer is a previously unobserved 6g/5a variation of the (6/5)AL motif and experienced a mix of both strong depletion and enrichment, often against the same target in replicate selection trajectories. Its coenrichment may therefore reflect the stochastic nature of the selection rather than a true positive coenrichment signal. Of the three aptamers of unknown structures that inhibit HXB2, only aptamer 80N 44 inhibits R277K. As expected, the F1Pk aptamers uniformly failed to inhibit.

When the 18 aptamers that inhibited both HXB2 and R277K were evaluated for inhibition of the panel of RTs used in the selection (**Figure 3B** and Figure 4), the (6/5)AL and UCAA motif aptamers were consistently broad-spectrum. Among the (6/5)AL aptamers, 70N 89 exhibited robust broad-spectrum inhibition of DDDP activity across the primate lentiviral RT panel; to a slightly lesser degree aptamers 70N 653 and 88.1 also inhibited broadly across the panel. These three aptamers all carry 6b/5b motifs. Aptamer A9, which also carries a 6b/5b motif, showed less inhibition and greater variability across the panel, potentially due to non-productive contributions of sequences surrounding the motif. Aptamer 70N 105, which carries a non-circularly permuted form of the 6b/5b motif, also exhibited a moderate to strong inhibition profile, as did the 6c/5b variant (80N 42A), which is a single nucleotide change from the 6b/5b motif. Other variations of the (6/5)AL motifs showed no discernible pattern of inhibition across the panel. In particular, 80N 433, which contains both a (6/5)AL and a UCAA motif, behaved most like ×1.1 in that it inhibited RTs from 94CY, MVP and EHO. RT from 94CY and EHO were inhibited more or less uniformly by all (6/5)AL aptamers tested, but inhibition was more variable for RTs from 93TH, MVP and TAN1B. Aptamers that carry UCAA motifs inhibited RTs from 94CY and EHO and exhibited mixed behavior against RTs from 93TH and MVP. Aptamer 80N 148 moderately inhibited RT from 93TH, but it failed against RT from MVP, while aptamer 80N 16 demonstrated the opposite pattern. All three UCAA aptamers inhibited RT from HIV-2 (EHO) to an extent that is comparable to that observed for the (6/5)AL aptamers, but they all failed to inhibit RT from TAN1B. The unknown aptamer, 80N 44, was only moderately broad-spectrum, inhibiting RTs from 94CY and EHO RTs, but none of the other RTs from non-subtype B lentiviruses. 3’ boundary determination and truncation experiments with 80N 44 against HXB2 RT identify a minimized 54 nucleotide sequence predicted to fold as a stem, interrupted with bulges and a (4/2) asymmetric loop, and capped with a six nucleotide loop (**Supplementary Figure S16**). The control F1Pk aptamers weakly inhibited RT from EHO, which is the only RT outside the HIV-1 subtype B to contain R277, but they failed to inhibit RTs from the other non-subtype B lentiviruses (**Figure 3B**).

**Figure 4.**
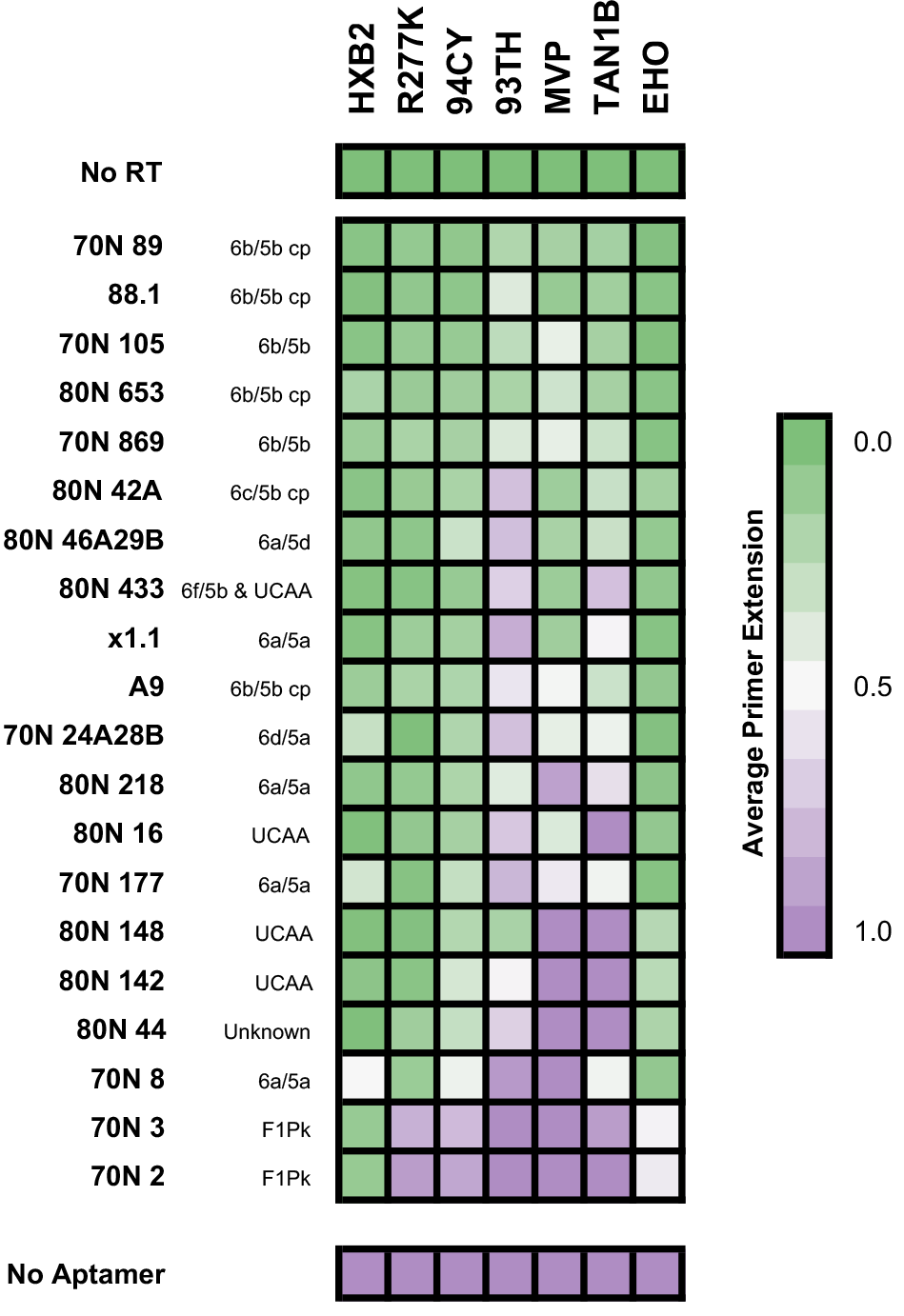
Inhibition profiles reveal motif-dependent broad-spectrum inhibition. Aptamers are sorted top-to-bottom by decreasing average inhibition, as measured by primer extension, against the panel of RTs used in the selection. Sorting in this manner reveals a pattern of biochemical RT inhibition corresponding to motif identity. (6/5)AL motifs, particularly the 6b/5b variant, demonstrated the strongest average inhibition against the tested RTs. UCAA motifs and the one previously-unidentified aptamer showed variable RT inhibition outside of Group M, and F1Pks showed no RT inhibition outside of Group M:B.

Sorting the aptamers by their average RT inhibition reveals a clear correlation of inhibition profiles with secondary structural motifs (**Figure 4**). Taken together, these data indicate that the (6/5)AL and UCAA motifs are capable of inhibiting a broad collection of non-subtype B RTs, and that several 6b/5b variants are capable of robust inhibition against every RT in our selection panel. While the UCAA and unknown motifs show limited inhibition outside of group M, they are potent inhibitors of the HIV-1 subtype A and B RTs tested here.

### Conversion of (6/5)AL variants to the canonical 6b/5b improves RT inhibition

Aptamers 88.1 (6b/5b), 80N 42A (6c/5b), and 80N 433 (6f/5b) all carry 6b/5b motifs or single-nucleotide variations, and they represent a gradient of inhibitory potency against RTs from non-B clades. To assess the relative contributions of nucleotide composition within and outside the asymmetric loop, we converted aptamer 88.1 from 6b/5b to 6f/5b and 6c/5b (**Figure 5A**) and each of the other two into canonical 6b/5b forms. These modifications are single nucleotide changes to interconvert AGCGUC (6b) to either AGCGU<u>A</u> (6f) or A<u>A</u>CGUC (6c), or vice versa. Establishing a canonical 6b/5b structural context for 80N 433 also required an A to G substitution immediately upstream of the 6b element to stabilize the flanking stem. The three original aptamers and the four variants were assayed for inhibition of RTs from HXB2, R277K, and five non-B clades (**Figure 5B**). Conversion of 88.1 away from the 6b/5b variant had little effect on the overall inhibition profile, with the exceptions of partial loss of inhibition for RTs from 93TH and TAN1B. In contrast, conversion of 80N 433 to 6b/5b (80N 433bb) dramatically increased its inhibitory potency, especially for RTs from 93TH, MVP and TAN1B, making 80N 433bb the strongest broad-spectrum inhibitor among all aptamers examined in this study. Conversion of 80N 42A to 6b/5b also improved its performance against the panel, with lowered mean primer extension values, albeit to a lesser degree than for 80N 433. Taken together, these data suggest that the exact composition of the 6 nucleotide bulge of the (6/5)AL loop is important for broad spectrum binding, that flanking sequences contribute directly or indirectly to net inhibition, and that the terminal C of the 6 nucleotide bulge, present in both 6b and 6c variants, may be especially important for establishing productive molecular contacts, particularly for RTs from 93TH and TAN1B.

**Figure 5.**
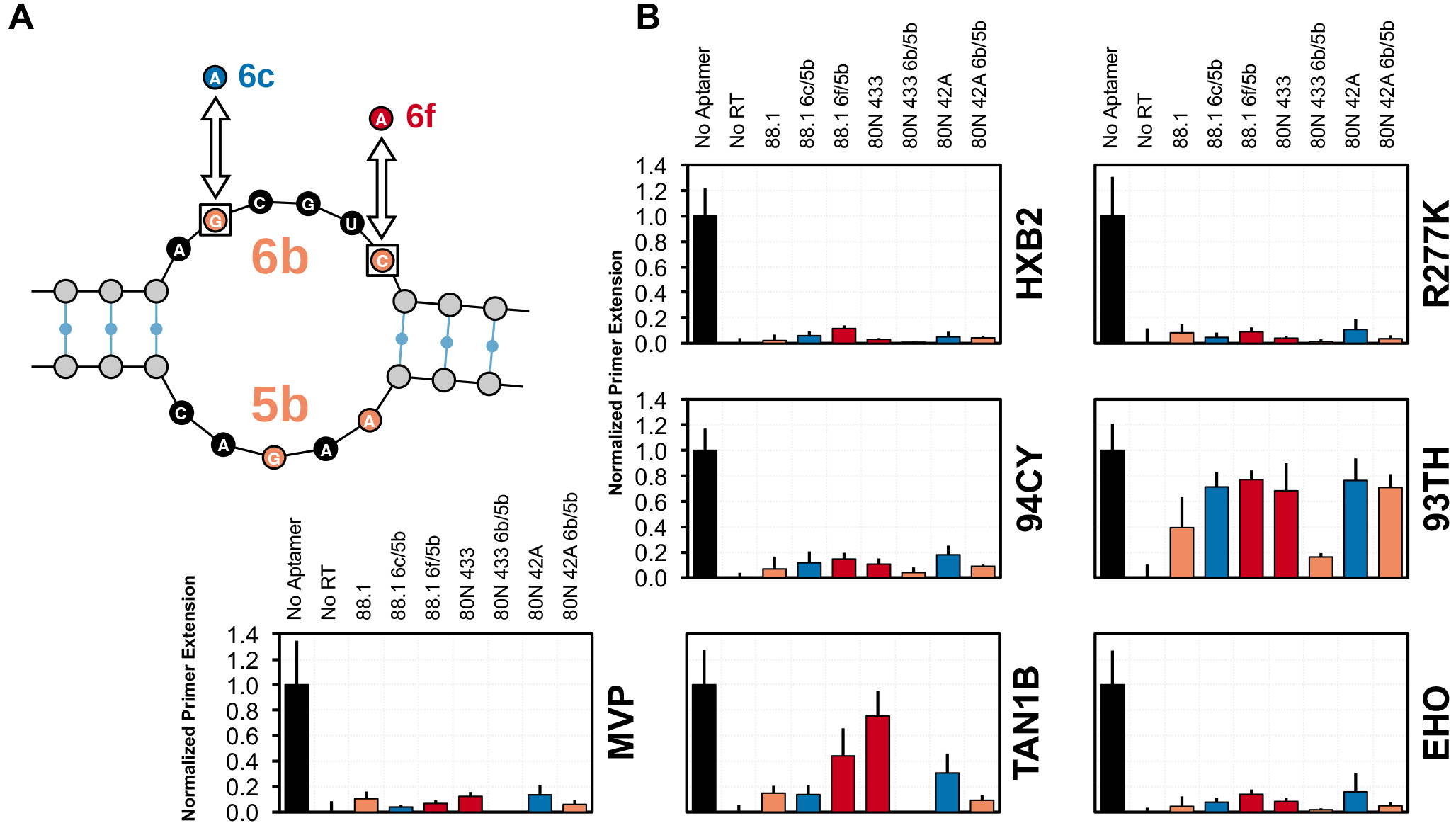
Conversion of exemplary (6/5)AL aptamers to the 6b/5b motif improves their broad-spectrum biochemical inhibition profile. (**A**) Single nucleotide conversions separate 6c/5b (blue) and 6f/5b (red) motifs from the 6b/5b motif (orange). Aptamer 88.1, which contains the 6b/5b motif, was converted to a 6f/5b or 6c/5b and assayed for changes in inhibition against RTs in the selection panel. Conversely, 80N 433 (a 6f/5b aptamer) and 80N 42A (a 6c/5b aptamer) were converted to 6b/5b. (**B**) Conversion towards the 6b/5b motif uniformly improved biochemical inhibition, whereas conversion away reduced inhibition in most cases. Mean values are shown (n ≥ 5) and error bars indicate standard deviation.

### Broad-Spectrum Aptamers Inhibit Outside of the Selection Panel

We reasoned that aptamers capable of inhibiting phylogenetically diverse primate lentiviral RTs likely interact with conserved features that would also be present in RTs that were not included in the PolyTarget selection panel. Anti-RT aptamers with broad-spectrum ability are therefore expected to bind and inhibit RTs that were not part of this “training set.” The aptamers chosen for this analysis were the most potent representatives of the three structural motifs that inhibited outside of subtype B: the 6b/5b converted form of aptamer 80N 433 (80N 433bb), a UCAA aptamer (80N 148) and the putative (4/2)AL motif (80N 44). The RTs for this assay sample two additional subtypes from HIV-1 group M and one from SIV_CPZ_. The group M RTs included a subtype C (98CN) and a circulating A/D recombinant form (92UG) for which the *pol* gene encoding RT groups within subtype D. The SIV_CPZ_ RT is from the US strain, derived from the chimpanzee species *Pan troglodytes troglodytes* (SIV_CPZ_*Ptt*), and is more evolutionarily related to HIV-1 group M subtypes than the TAN1B SIV strain used in the selection panel (SIV_CPZ_*Pts*) (**Supplementary Figure S1**) (23). Against the RT from subtype D (92UG), the UCAA aptamer showed moderate inhibition and the 6b/5b aptamer showed complete inhibition (**Figure 6**). When assayed against the RT from subtype C (98CN), all three aptamers showed moderate inhibition, with the 6b/5b aptamer exhibiting virtually complete inhibition of the DDDP activity. Finally, only the 6b/5b aptamer inhibited the RT from SIV_CPZ_ (US). These results provide further evidence of the ability of the UCAA and unknown motif to inhibit non-subtype B RTs and of the 6b/5b motif to strongly inhibit RT from phylogenetically diverse primate lentiviral strains.

**Figure 6.**
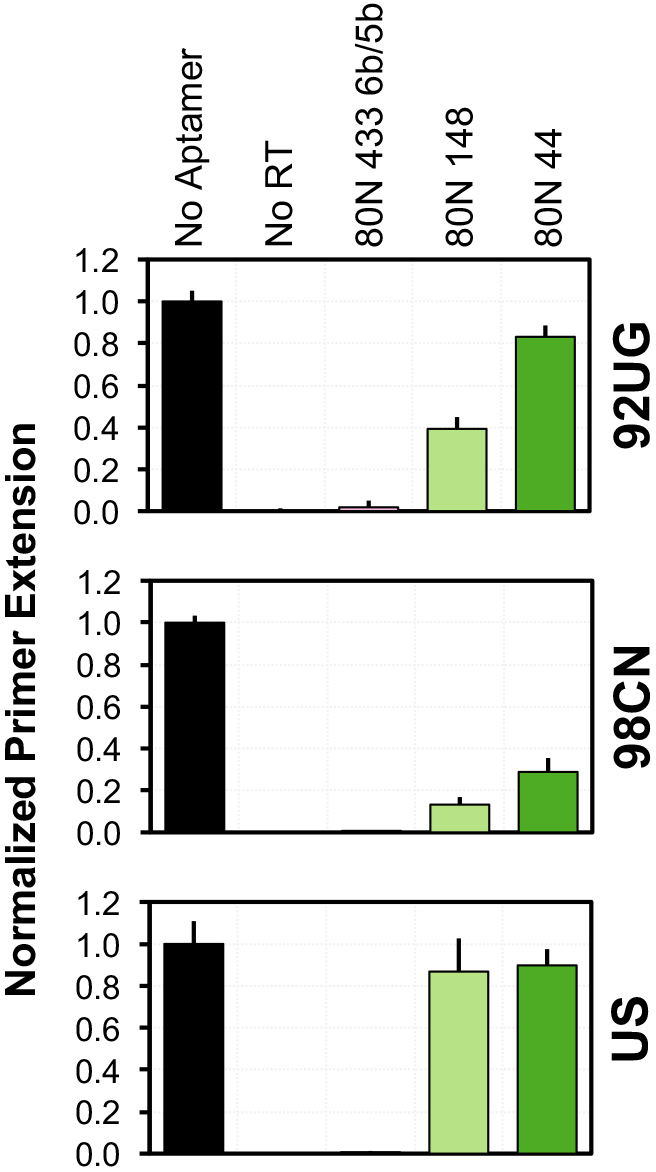
Broad-spectrum aptamers are capable of RT inhibition outside of the selection panel. Aptamers representing the three motifs with non-subtype B RT inhibition were tested against RTs from two group M strains (92UG and 98CN) and an SIV strain (US). 92UG, a circulating recombinant form of HIV-1 group M whose RT groups with subtype D, is inhibited moderately by the UCAA motif aptamer and robustly by the 6b/5b aptamer. RT from 98CN, a group M:C strain, is inhibited by all three aptamers tested, including 80N 44 (unknown motif). RT from the US strain of SIV_CPZ_*Ptt*, is only inhibited by the 6b/5b aptamer. Mean values are shown (n ≥ 5) and error bars indicate standard deviation.

### Broad-Spectrum Anti-RT Aptamers Bind with Low Nanomolar Affinity

We hypothesized that the observed patterns of sequence enrichment and enzymatic inhibition above should reflect relative affinities for each aptamer-RT combination. For example, because modifying aptamer 80N 433 into a canonical 6b/5b form improved its inhibition of RTs from 93TH and TAN1B RT, the modified aptamer is expected to exhibit improved affinity for these RTs. Apparent dissociation constants for a number of aptamer-RT pairs were determined by a nitrocellulose filter binding assay (**Table 1 and Supplementary Figure S17**). As expected, aptamer 70N 2 (F1Pk) bound well to RT from HXB2 (Kd ≈ 110 nM) but showed no discernable affinity for the R277K point mutant, consistent with its inhibition patterns (**Figure 3A**). Furthermore, its affinity for RT from HXB2 was notably weaker than affinities observed for the broad-spectrum aptamers tested here, consistent with the moderate depletion of aptamer 70N 2 in the HXB2 trajectory and its powerful depletion in each of the others (**Figure 2 and Supplementary Table S5**). Dissociation constants for all of the other aptamers were in the low nanomolar range (< 100 nM) for binding to HXB2. Because these assays were all performed at the same time using the same protein preparation, we conclude that the affinity of aptamer 80N 433bb for RT from HXB2 is 5-fold improved relative to that of the pseudoknot aptamer 70N 2. All of the broad-spectrum aptamers showed reduced affinity for the R277K mutant, relative to their affinities for RT from HXB2. For the RT’s from non-subtype B lentiviruses, relative binding affinities among the broad-spectrum aptamers correlated with their inhibition profiles. In particular, Kd values for the 6b/5b variant of 80N 433 were in the low nanomolar range across the RT panel, and this aptamer uniformly bound more strongly to each RT than did any of the other aptamers. Affinity improvements associated with converting the original 80N 433 from 6f/5b to 6b/5b ranged from approximately 3-fold for HXB2 and 93TH to 13-fold for TAN1B, illustrating the variable contributions of these nucleotide changes upon the RT-aptamer interactions.

**Table 1.** **Apparent binding affinity (*K*_*d*_) of broad-spectrum RT aptamer-inhibitors^a^**

|  | **HXB2** | **R277K** | **94CY** | **93TH** | **MVP** | **TAN1B** | **EHO** |
| --- | --- | --- | --- | --- | --- | --- | --- |
| **70N 2** | 110 ± 20 | >>300 | - | - | - | - | - |
| **80N 44** | 60 ± 12 | 190 ± 51 | 30 ± 9 | 58 ± 17 | - | - | - |
| **80N 148** | 74 ± 17 | 150 ± 42 | 55 ± 13 | 13 ± 3 | - | - | - |
| **80N 433** | 59 ± 23 | - | - | 22 ± 6 | - | 210 ± 38 | - |
| **80N 433bb** | 22 ± 4 | 38 ± 7 | 29 ± 8 | 7 ± 1 | 39 ± 8 | 16 ± 2 | 34 ± 9 |

<sup>a</sup>Values are listed in nanomolar with standard deviation from triplicate assays. Binding curves are provided in Supplementary Figure S17.

### Broad-Spectrum Anti-RT Aptamers Suppress Viral Replication in Cell Culture Assays

To determine whether aptamers that demonstrate broad-spectrum activity *in vitro* also inhibit diverse HIV in a biological context, we adapted an assay that we had previously developed for monitoring aptamer-mediated inhibition of HIV replication in cell culture (12, 13, 27). Briefly, transfection of plasmids encoding a replication-deficient HIV reporter virus and the glycoprotein from vesicular stomatitis virus (VSV-G) leads to production of virus that are competent for a single cycle of replication and for which transfection and infection efficiencies can both be measured using the virally-encoded EGFP. Cotransfection of aptamer-expressing plasmid simultaneously with viral plasmids can lead to packaging of the aptamer into the virus during assembly, inhibition of reverse transcription in the target cells, and loss of infectivity that can be measured by a loss of fluorescence signal in the target cells. To incorporate phylogenetically diverse RT subtypes into this assay, we first built and evaluated proviral plasmids to express several RTs from our Poly-Target selection panel (HXB2, R277K, 93TH, 94CY, and MVP) such that only the RT segments differed in each construct (13). This assay was used to evaluate viral inhibition by aptamers that demonstrated broad-spectrum RT inhibition *in vitro* by cotransfecting various combinations of proviral and helper VSV-G plasmids with aptamer or control plasmids, collecting the pseudotyped virus, and determining the percent of EGFP-positive cells after infection of fresh 293FT cells (**Figure 7**). Infectivity was normalized to p24 levels determined by ELISA. As observed previously (13), aptamer 70.05 (an F1Pk pseudoknot) strongly inhibited infectivity of virus carrying the HXB2 RT, but not the R277K point mutant or any of the non-subtype B RTs. In contrast, aptamers 80N 433 and 80N 433bb inhibited infectivity of all recombinant viruses in the panel. Aptamers 80N 148 and 80N 44 demonstrated a variable pattern against the non-subtype B RTs. Both inhibited replication by viruses carrying RT from 94CY but only 80N 148 inhibited virus carrying RT from 93TH, even though the RTs from 94CY and 93TH both group within subtype A and are 91.4% identical (98.2% similar). Neither 80.148 nor 80.44 inhibited viruses carrying RT from the MVP5180 strain of HIV-1 Group O. These results are consistent with our biochemical inhibition results and closely mirror our previous observations for inhibition by the major anti-RT motifs in cell culture (13).

**Figure 7.**
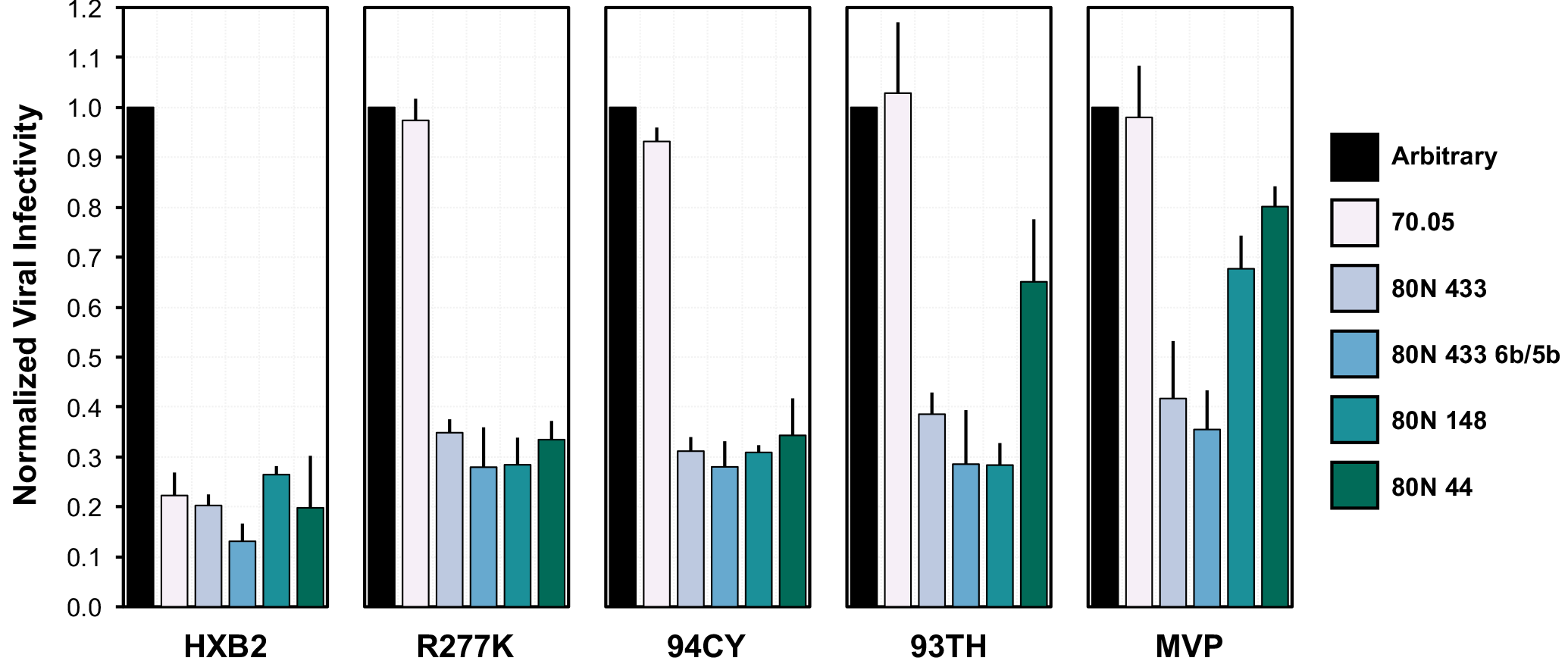
Broad-spectrum anti-RT aptamers suppress viral replication in biological assays. A non-inhibitory RNA (“arbitrary”), a previously characterized F1Pk aptamer (70.05), and several broad-spectrum aptamers identified through Poly-Target selection were assessed for their ability to suppress virus in a single-cycle infectivity assay.

## DISCUSSION

Recent advances in aptamer technology have enabled increased selection throughput and high-resolution analysis of selection outcomes (28, 29). Microfluidic (30) and multiplexed selection platforms (31, 32) facilitate efficient and simultaneous selection against numerous targets, and emulsion methods that encapsulate single genotypes into “monoclonal aptamer particles” are readily partitioned or sorted according to phenotype (33, 34). Post-selection, high-throughput sequencing (HTS) has empowered optimization of candidate aptamers based on round-to-round enrichment of sequences and by providing large data sets that allow for powerful bioinformatic analyses of sequence, structure, and function relationships (7, 35–37). In addition to advances in selection methodology and candidate identification, high-throughput aptamer characterization platforms that utilize microarrays can screen thousands of aptamers simultaneously (38), and can directly link genotype to phenotype when performed on a sequencing platform (39, 40). Here we sought to combine HTS with multiplexed selections against a diverse panel of recombinant RTs. We applied these technologies to elicit specialized binding and inhibition phenotypes (cross-reactivity) from well-characterized and pre-enriched aptamer libraries. Furthermore, we sequenced negative selection populations and performed our positive selections in duplicate. The combination enabled coenrichment analysis of multiple selection trajectories, nuanced with data from duplicate and negative selections. This strategy efficiently narrowed the number of potential candidates to an examinable amount, allowing us to identify highly-potent RNA broad-spectrum inhibitors of primate lentiviral RTs among these candidates.

Other selection approaches have previously been used to identify cross-reactive aptamers or to exploit branched selections. In “toggle SELEX” (41), the target of a single-selection trajectory is alternated, or “toggled” between rounds; this approach was used to identify inhibitors of human and porcine thrombin and later to identify broadly-reactive aptamers targeting whole bacterial cells (42). In another approach, single-pot selections against numerous targets were used to identify cross-reactive aptamers against several M-types of *Streptococcus pyogenes* (43) and against HA proteins from multiple strains of Influenza A (44). Although both of these methods are powerful (alternating or simultaneous targets), they rely on a single evolutionary trajectory and lack the target-specific information that can inform the aptamer selection and design process. Recently, Dupont *et al.* reported on a clever branched selection approach against alanine point mutants of serpin plasminogen activator inhibitor-1 (PAI-1) (45). Utilizing a library previously selected against wild type pAI-1, those authors performed one additional round of selection against a panel of 10 targets, in parallel, followed by high-throughput sequencing, to identify aptamer binding site preferences. Unlike the Poly-Target selection described here, the pre-enriched library utilized by Dupont *et al.* had not been saturated. In the original selection against PAI-1, enrichment of the library was detected after 5 rounds and proceeded for a total of 8 rounds (46). Rather than using the heavily-enriched library from round 8, the branched selection began with round 5, allowing for a greater diversity of aptamers to emerge.

The Poly-Target approach utilized pre-enriched libraries to simultaneously explore a wider sequence space than available in a traditional aptamer directed evolution strategy. Interestingly, neither abundance, nor enrichment, nor coenrichment alone was a strong predictor of fitness in the Poly-Target selection performed here. Highly abundant sequences in the starting libraries depleted across most selection trajectories yet retained a major presence within the Round 17 populations. In contrast, most of the 6b/5b variants that enriched through the selection were so rare in the starting populations that, despite their large fold-enrichment, they remained as a relatively small percentage of the populations and could only be reliably identified through HTS. Coenrichment was slightly more predictive of broad-spectrum phenotypes after removing aptamers that enriched against nitrocellulose alone. Nevertheless, several F1Pk aptamers demonstrated enrichment across several trajectories despite their lack of broad-spectrum inhibition. The combined use of coenrichment and codepletion analysis of sequence clusters identified sequences that experienced mixed enrichment and depletion, either across trajectories or between replicates. The most productive analytic strategy for the present datasets combined each of these approaches with careful curation based on sequence and structural motifs established from previous investigations. The Poly-Target approach may be broadly applicable for any combinatorial selection or directed evolution strategy that seeks to identify broad-spectrum behavior given an appropriate phenotypic screen.

The primer-template-binding groove in RT extends from between the thumb and fingers domain along the palm to the RNaseH domain. Although the RT amino acid sequence within this groove is among the most highly-conserved portions of the protein, natural polymorphisms and subtype-specific variations affect nucleic acid interaction affinity and specificity. The Poly-Target selection strategy was designed to intentionally select for promiscuity against a family of related targets, rather than specificity for a single target or its point-mutants, and it relied on the sequencing depth provided by HTS to identify functionally rare aptamers that enriched from our Round 14 starting libraries. We developed and utilized a coenrichment analysis strategy to efficiently reduce the dataset from over a hundred million sequences to focus on dozens of candidate sequence clusters for further analysis. We were able to identify several rare variants of (6/5)AL motifs and also a previously-unknown aptamer inhibitor of HIV-1 group M RTs. Inhibition assays identified the 6b/5b variant of the (6/5)AL motif as an especially potent inhibitor of the DDDP activity of RTs utilized in the selection panel and several RTs that were not represented in the selection. Conversion of aptamer 80N 433 towards the 6b/5b motif improved its inhibition profile across the panel of RTs and most notably against 93TH, MVP and TAN1B. Furthermore, the UCAA aptamer (80N 148) and aptamer 80N 44 demonstrated inhibition of RT outside of subtype B strains. Follow-up assays correlated *in vitro* inhibition of RTs with binding affinity and suppression of viral replication. As our efforts were focused on identification of broad-spectrum inhibitors, we did not attempt to find strain-specific inhibitors nor did we closely examine the role of individual nucleotides and their interactions with specific RTs. The wealth of data provided by the PolyTarget selection approach should, in principle, allow one to look for enrichment exclusive for a particular target, in addition to advancing the broad-spectrum objectives that were the focus of the present study. An intriguing question to be explored in future studies is whether resistance arises from a small number of mutations for ‘specialist’ aptamers (such as the R277K mutation for F1Pk aptamers) and requires significantly more mutations for broad-spectrum aptamers, making such breakout mutation a low-probability event and creating what is sometimes referred to as a high genetic barrier to resistance.

## MATERIAL AND METHODS

### RT Expression and Purification

DNA encoding the p66 and p51 subunits of each RT (**Supplementary Table S1**) were PCR amplified from previously described pET200/D-TOPO expression vectors (16). Amplicons were then restriction digest cloned into the pRT-Dual plasmid, a derivative of pETDuet-1 vector (a gift from Stefan Sarafianos (47)). Plasmids expressing RTs from HXB2, R277K, 94CY and TAN1B utilized the PpuM1/SacI cloning sites for the p51 subunit, and the Sacll/Avrll sites for the p66 subunit (“pRTD” series plasmids). Whereas RTs from 93TH, MVP and EHO utilized the same restriction sites for the p66 subunit but utilized Notl/Sall for the p51 subunit (“pRTD2” series plasmids). Additional plasmid engineering was performed on the ribosome binding site upstream of the p66 subunit in the pRTD2 series plasmids to optimize expression. Plasmid inserts were confirmed by Sanger sequencing (University of Missouri DNA Core Facility) and are available upon request.

RT expression plasmids were heat shock transformed into *E. coli* BL21(DE3)pLysS competent cells and then incubated overnight at 37° C on LB-Agar plates supplemented with selective antibiotic (50 μg/mL streptomycin). Clonal isolates from each plate were recovered and incubated for approximately 16 hours at 37° C with 250 RPM shaking in 10 mL of 2xYT selective liquid media. Cultures were then diluted into 1 L of fresh media and optical density at 600 nm was monitored until reaching 0.5, whereupon protein overexpression was induced with 1 mM IPTG. Following a 4-hour incubation, samples were centrifuged for 15 minutes at 4,000 RCF. Supernatants were decanted and cell pellets were frozen at −80° C until purification. For purification, cell pellets were resuspended in sonication buffer (25 mM Tris-HCl pH 8.0, 500 mM NaCl, 1 mM PMSF, 0.15 mg/mL lysozyme) and then subjected to four ultrasonication steps for 30 seconds on ice, with 2 minutes of rest on ice in between steps. Cell lysates were then centrifuged for 15 minutes at 4° C and 12,000 RCF to remove cell debris. Cell extracts were then passed through a 0.45 μm filter to remove insoluble material and applied to a Ni-NTA agarose affinity column (Qiagen) for His tag purification. Purification was performed according to the manufacturer’s protocol with an added high salt wash (2 M NaCl) to remove any endogenous nucleic acids bound to RTs. Eluted protein samples were pooled and quantified using UV absorbance at 280 nm on a NanoDrop 1000 spectrophotometer (Thermo Fisher) with estimated extinction coefficient and molecular weight information. Purified proteins were concentrated using Amicon Ultra Centrifugal Filters (Millipore Sigma) and dialyzed into 2× storage buffer (100 mM HEPES pH 7.5, 100 mM NaCl) using Slide-A-Lyzer cassettes (Thermo Scientific). RT preps were validated for purity and size by SDS-PAGE, and for activity by comparison of primer extension assays against previous preps. Proteins were stored at −20° C after addition of glycerol to 50% (v/v).

### Poly-Target Selection

Starting libraries for this work were the Round14 70N and 80N libraries described previously (6). Double-stranded DNAs from these libraries were transcribed using T7 RNA polymerase, *in vitro* transcription buffer (50 mM Tris-HCl pH 7.5, 15 mM MgCl_2_, 5 mM DTT, and 2 mM spermidine), and 2 mM of each ATP, CTP, GTP and UTP. Reactions were incubated at 37° C for a minimum of 4 hours and halted with the addition of gel loading buffer (95% formamide, 5mM EDTA, xylene cyanol FF and bromophenol blue). Transcribed RNAs were purified through denaturing polyacrylamide gel electrophoresis (6% TBE-PAGE, 8 M urea) and bands corresponding to the expected size were gel extracted and eluted while tumbling overnight in 300 mM Sodium Acetate pH 5.4. Eluates were ethanol precipitated, resuspended in TE buffer (10 mM Tris-HCl pH 8.0, 1 mM EDTA), and stored at −20° C until use. RNA concentrations were determined on a NanoDrop 1000 spectrophotometer (Thermo Fisher). For each trajectory and round of selection, 200 pmol of the transcribed libraries were resuspended in 100 μL binding buffer (150 mM KCl, 10 mM MgCl_2_, 50 mM Tris-HCl pH 7.5) and renatured by heating to 65 °C and cooling on ice. 40 pmol of the respective RT was then added, and the mixture was incubated on ice for an additional 20 minutes. Separately, a 25 mm nitrocellulose filter (#HAWP02500, Millipore Sigma) was pre-wet with 2 mL binding buffer on a sampling manifold (#XX2702550, Millipore Sigma). Immediately before application of incubated RNA:RT complex, the nitrocellulose filter was washed again with 1 mL of binding buffer under applied vacuum. RNA:RT complex was then applied to filter under vacuum and washed with 1 mL of binding buffer. Suction continued for 10 minutes. The filter was then removed and incubated in extraction buffer (8 M urea, 50 mM NaCl, 10 mM EDTA). RNA was recovered by phenol:chloroform extraction of the filter and ethanol precipitation. Recovered RNA was reverse transcribed using ImProm-II Reverse Transcriptase (Promega) and PCR amplified to repeat the selection process, or for high-throughput sequencing (see below). Prior to Round 15, transcribed Round 14 libraries were incubated with a thin strip of nitrocellulose for several minutes to subtract non-specific binders and RNA was extracted and purified as described above, with the exclusion of the RT binding and partitioning steps. Nitrocellulose-binding RNAs were recovered and reverse transcribed for HTS, while nitrocellulose-subtracted RNAs were used for independent selections.

### High-throughput Sequencing and Bioinformatics Analysis

Libraries were prepared for sequencing using a series of PCR steps to add Illumina adapters and sequencing indices for multiplexing of the 70N and 80N libraries as previously described (7). Sequencing was performed on an Illumina HiSeq2000 (University of Missouri DNA Core Facility). Populations were demultiplexed to identify and parse the 5’ and 3’ constant regions. Data preprocessing was performed using cutadapt (48) to trim 5’ and 3’ constant regions from sequences and to discard any uncut sequences or sequences not within ± 3 of the expected size (70 or 80) after trimming. Trimmed sequences were then filtered for high-quality reads using FASTQ quality filter from the FASTX-Toolkit (http://hannonlab.cshl.edu/fastx_toolkit/). Quality filtering eliminated a sequence if a single position had a Phred quality score of less than 20. Trimmed and quality filtered sequences were then processed using the FASTAptamer toolkit (24) to count and normalize sequence reads (FASTAptamer-Count), calculate fold-enrichment from Round 14 to Round 17 (FASTAptamer-Enrich), compare sequence frequencies across populations (FASTAptamer-Compare), group related sequences into clusters (FASTAptamer-Cluster) and search for known sequence motifs (FASTAptamer-Search). Custom scripts (open source and available at https://github.com/FASTAptamer/PolyTarget) were written in Perl to perform coenrichment analysis, to map sequences onto their Round 14 identity, and to recluster the sequences as described in the results. RNA secondary structures were predicted using the mfold webserver (26) and depicted using VARNA: Visualization Applet for RNA (49).

### Aptamer Generation

Oligonucleotides for DNA transcription templates were ordered from Integrated DNA Technologies and ligated as previously described (50). Ligated oligonucleotides were then PCR amplified with Pfu DNA polymerase and primers containing the T7 promoter and the remainder of the constant region that was not included in the oligonucleotides. Amplification products were verified for size using agarose gel electrophoresis. Run-off transcription reactions using T7 RNA polymerase and RNA extraction and purification were performed as described above for transcription of Poly-Target libraries. RNAs were refolded by heating to 65° C and cooling on ice prior to use.

### Primer Extension Assays

DNA-dependent DNA polymerase activities of RTs were assayed using a 31 nucleotide DNA template (5’ CCATAGATAGCATTGGTGCTCGAACAGTGAC 3’) and a complementary, 5’-Cy3-labeled 18-nucleotide DNA primer (5’ Cy3-GTCACTGTTCGAGCACCA 3’) as previously described (22). In short, 20 nM of RT and 100 nM of aptamer (omitted for “*No Aptamer*’ control) were pre-incubated on ice for 10 minutes in extension buffer (50 mM Tris-HCl pH 7.5, 50 mM NaCl, 5 mM MgCl_2_). 10 nM primer, 20 nM template and 100 μM of each dNTP were then added immediately before incubation at 37° C for 10 minutes. Reactions were halted with the addition of gel loading buffer (95% formamide, 5 mM EDTA, xylene cyanol FF and bromophenol blue) and analyzed by denaturing polyacrylamide gel electrophoresis (10% TBE-PAGE, 8 M urea). Gels were scanned on a Typhoon FLA9000 imager (GE Healthcare) and quantified using Multi Gauge software (Fujifilm) for fraction of primer extended. Primer extension values were then normalized by subtracting the mean “*No RT*’ value for the respective RT and multiplying by a normalization factor that defines 100% extension as the mean extension in the “*No Aptamer*’ control.

### Aptamer:RT 3’ Boundary Determination

*In vitro* transcribed and purified RNA was treated with Antarctic phosphatase (Fermentas) to remove the 5’ terminal phosphate and subsequently labeled with T4 polynucleotide kinase in the presence of Y-^32^P labeled ATP (Perkin Elmer). Radiolabeled RNA was gel purified by denaturing PAGE as described for transcription of libraries. RNase T1 digestion was performed by incubating thermally renatured RNA (> 10^6^ CPM) with 40 units of RNase T1 (Thermo Fisher) in digestion buffer (25 mM sodium citrate pH 5.0, 6 M urea) for 5 minutes at 55° C. T1 digestion was halted with the addition of gel loading buffer. Alkaline hydrolysis was performed by incubating RNA in 50 mM sodium carbonate pH 9.0 at 90° C for 10 minutes. Hydrolysis was halted upon addition of 300 mM sodium acetate pH 5.0 and were then ethanol precipitated and resuspended in H_2_O. Aptamer:RT 3’ boundary was determined by incubating 50 pmol RNA with 100 pmol HXB2 RT and partitioning the bound complexes as described above for the Poly-Target selection. RNAs were recovered from the filter and resolved on a denaturing polyacrylamide gel (15% TBE-PAGE, 8 M urea).

### Affinity Constant Determination

Affinity constants (*K*_*d*_’s) were determined by a radiolabelled binding assay. Approximately 15-20k CPM of labeled and refolded RNA was incubated with varying concentrations of RT (0.1 – 1000 nM, or without RT to determine background binding) in binding buffer (50 mM Tris-HCl pH 7.5, 140 mM KCl, 1 mM MgCl_2_, 0.1 μg/mL bovine serum albumin) and allowed to equilibrate on ice for 15 minutes. Assembled RNA:RT complexes were then partitioned from unbound RNA by passing samples over a nitrocellulose filter (#HAWP02500, Millipore Sigma) under vacuum (#XX2702550, Millipore Sigma) and immediately washing with 500 μL binding buffer. Radioactivity retained on filters was counted by adding 4 mL of scintillation fluid to filters placed inside of scintillation vials and counted using a liquid scintillation counter. Fraction of RNA bound was calculated by determining the fraction of radioactivity bound and were fit to a one site, specific binding curve using Prism GraphPad 6.2.

### Viral infectivity assay

Cell culture, virus production and evaluation of viral infectivity were carried out as previously described (13, 27). Single-cycle infectivity assays using aptamer-expressing plasmids, pNL4-3-CMV-GFP and pMD-G (VSV-G), were performed by transfecting 293FT cells with polyethylenimine (PEI) in six-well cell culture dishes. All transfections were performed on cells plated the previous day (50% confluence) using PEI at 3 μl/μg DNA, as previously described. Aptamer-expressing plasmids (1000 ng) were cotransfected with a mixture of pNL4-3-CMV-GFP (150 ng) and pMD-G (50 ng). The medium was changed approximately 12 hours after transfection. Viral supernatant was harvested 48 hours after the post-transfection media change and clarified by centrifugation to remove cellular debris. Cell-free viral supernatant (50 μl) was added to fresh 293FT cells to determine infectivity. Infected cells were collected 24-48 hours post-infection, fixed with 4% paraformaldehyde and analyzed on an Accuri C6 Flow Cytometer (BD Biosciences) to determine the percentage of infected (EGFP-positive) cells. Infectivity data was normalized to levels of p24 determined by ELISA.

## ACKNOWLEDGEMENTS

The authors would like to thank Katherine N. Wilsdon for technical assistance.

## Supplementary Data

**Figure S1.**
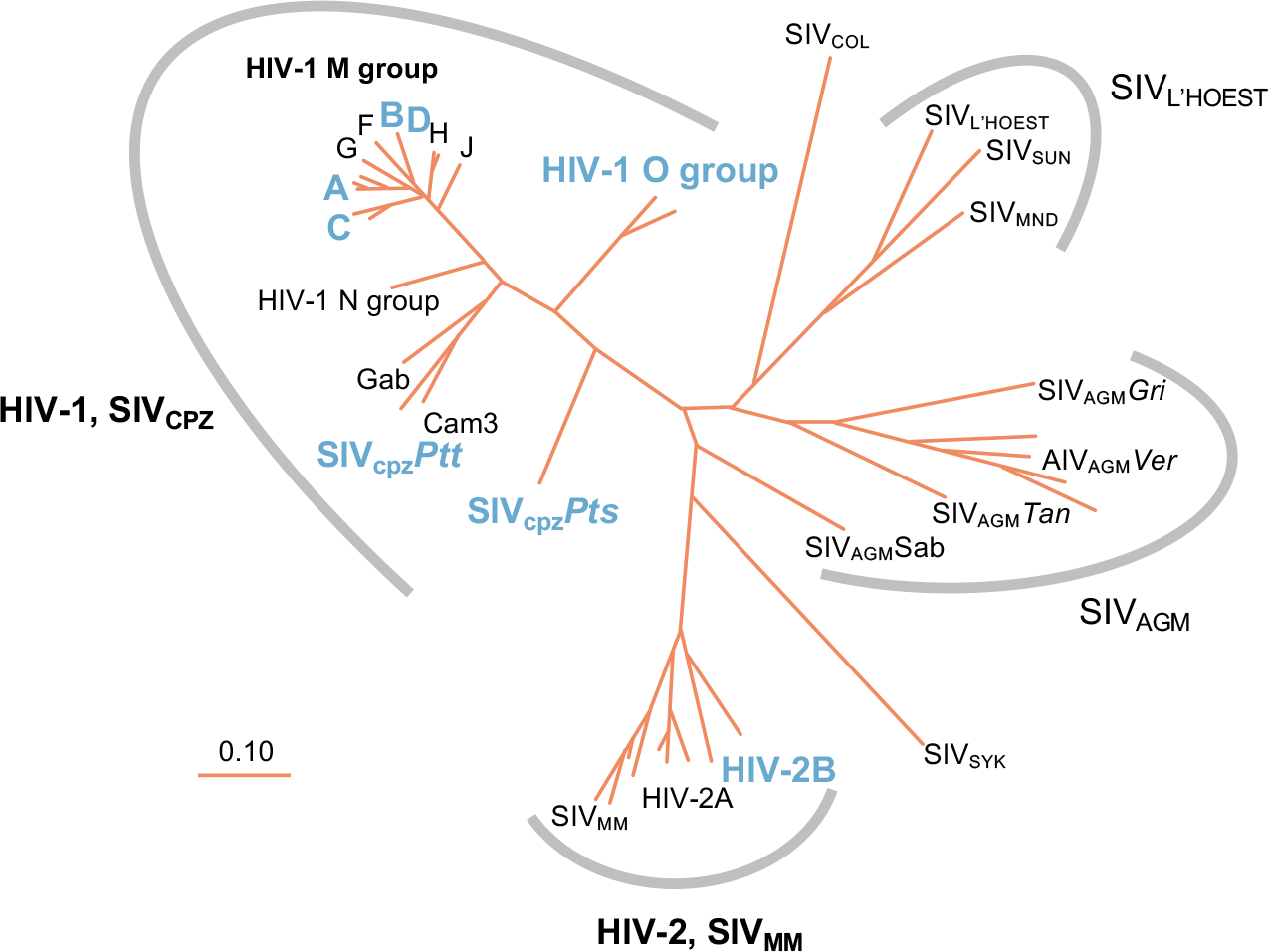
Phylogenetic relationships among the primate lentivirus *Pol* gene. The lentiviral *Pol* gene codes for reverse transcriptase, protease, RNase, and integrase. Relative to the rest of the viral genome, *Pol* is highly conserved and is therefore used to construct phylogenetic relationships among lentiviral strains. The phylogenetic tree depicted here illustrates the genetic diversity both within and across lentiviral groups. Reverse transcriptase from the groups and subtypes used in this study are shown in bold and colored blue. Adapted, with permission, from The Human Retroviruses and AIDS 1999 Compendium (https://www.hiv.lanl.gov/).

**Table S1.** Panel of recombinant RTs used in Poly-Target selection and characterization.

| Virus (Group:Subtype) | Strain<sup>a</sup> | GenBank Accession Number | Differences from Reference Sequence |
| --- | --- | --- | --- |
| HIV-1 (M:B) | **HXB2** | K03455 | - |
| HIV-1 (M:B) | HXB2 (**R277K**) | K03455 | R277K<sup>b</sup> |
| HIV-1 (M:A) | **94CY**017.41 | AF286237 | S554T |
| HIV-1 (M:CRF01_AE) | **93TH**253.3 | U51189 | D6E, I553S, S554T |
| HIV-1 (O) | **MVP**5180 | L20571 | M16V, I184M, V261I, V435I, N465K, E523K |
| SIV<sub>Cpz</sub>*Pts* | **TAN1B** | AF447763 | - |
| HIV-2 (B) | **EHO** | U27200 | E15G, K64R, K66R, N176S, K206R, A449V, E520K |
| HIV-1 (M:A/D)<sup>c</sup> | **92UG**021 | AF009396 | D121C, E169D, V293I |
| HIV-1 (M:C)<sup>c</sup> | **98CN**009 | AF286230 | R102K, S554T |
| SIV<sub>Cpz</sub>*Ptt*<sup>c</sup> | **US** | AF103818 | Y208H, E223K, H315R, R345Q, L392P, G498D, V518A |

<sup>a</sup>Abbreviated names (in bold) are used throughout document to refer to the strains listed here.
<sup>b</sup>R277K mutation only present in p66 subunit.
<sup>c</sup>Reverse transcriptases not included the selection panel, but used to characterize broad-spectrum inhibition.

**Figure S2.**
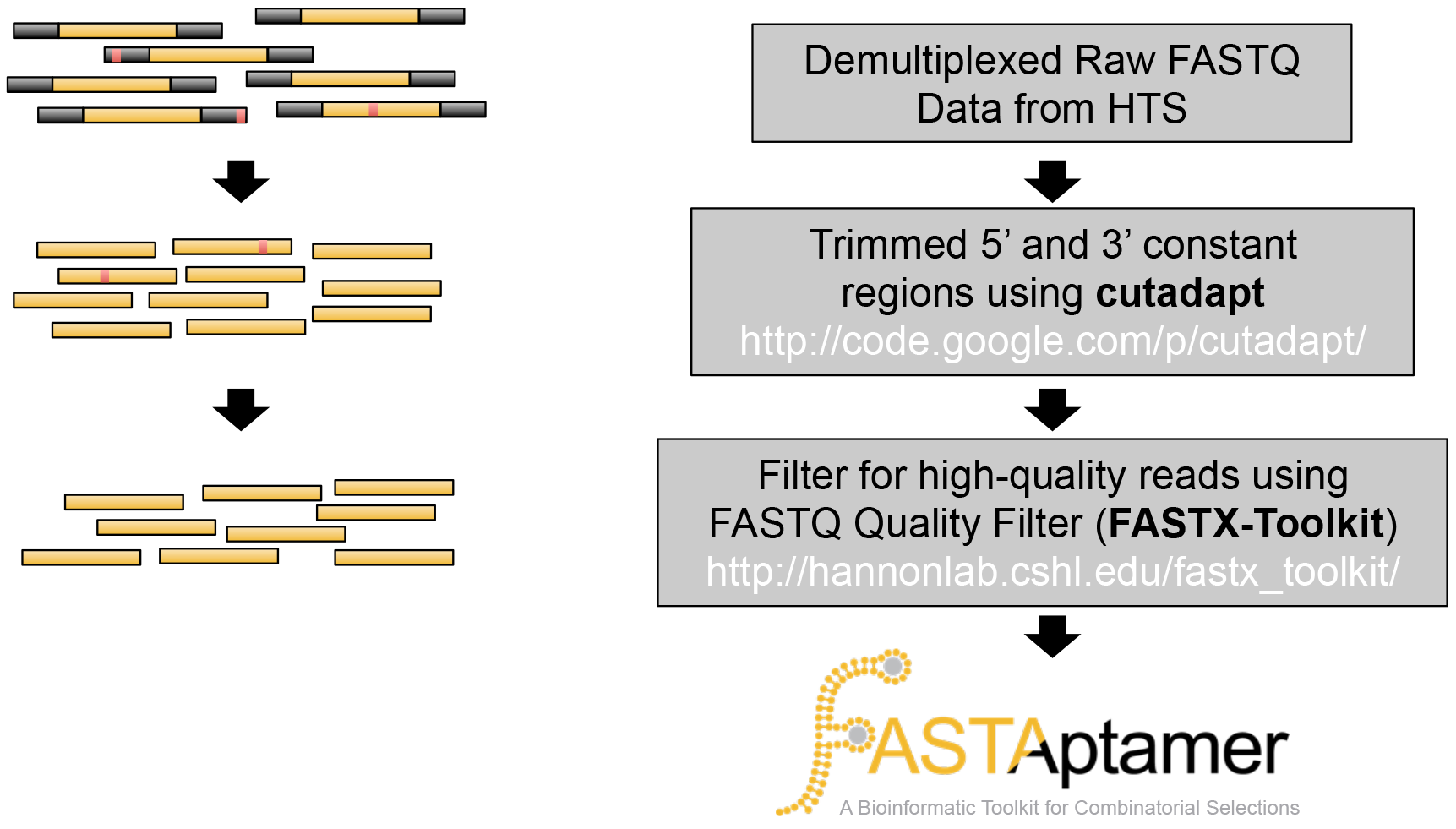
Pre-processing workflow for sequencing of Poly-Target selection populations. High-throughput sequencing populations were pre-processed to isolate only high-quality reads of the randomized regions. Raw FASTQ files were demultiplexed into individual populations using custom Perl scripts. Constant regions (represented by the black bars) were then removed using cutadapt (48). Trimmed reads were filtered using FASTQ Quality Filter from the FASTX-Toolkit (http://hannonlab.cshl.edu/fastx_toolkit/) to remove sequences that contained any nucleotides called with < 99% confidence (red bars). Trimmed and filtered files were then processed using the FASTAptamer toolkit to determine sequence frequencies, compare selection populations, calculate fold-enrichment from the Round 14 starting libraries, group related sequences into clusters and search for sequence motifs (24).

**Table S2.**
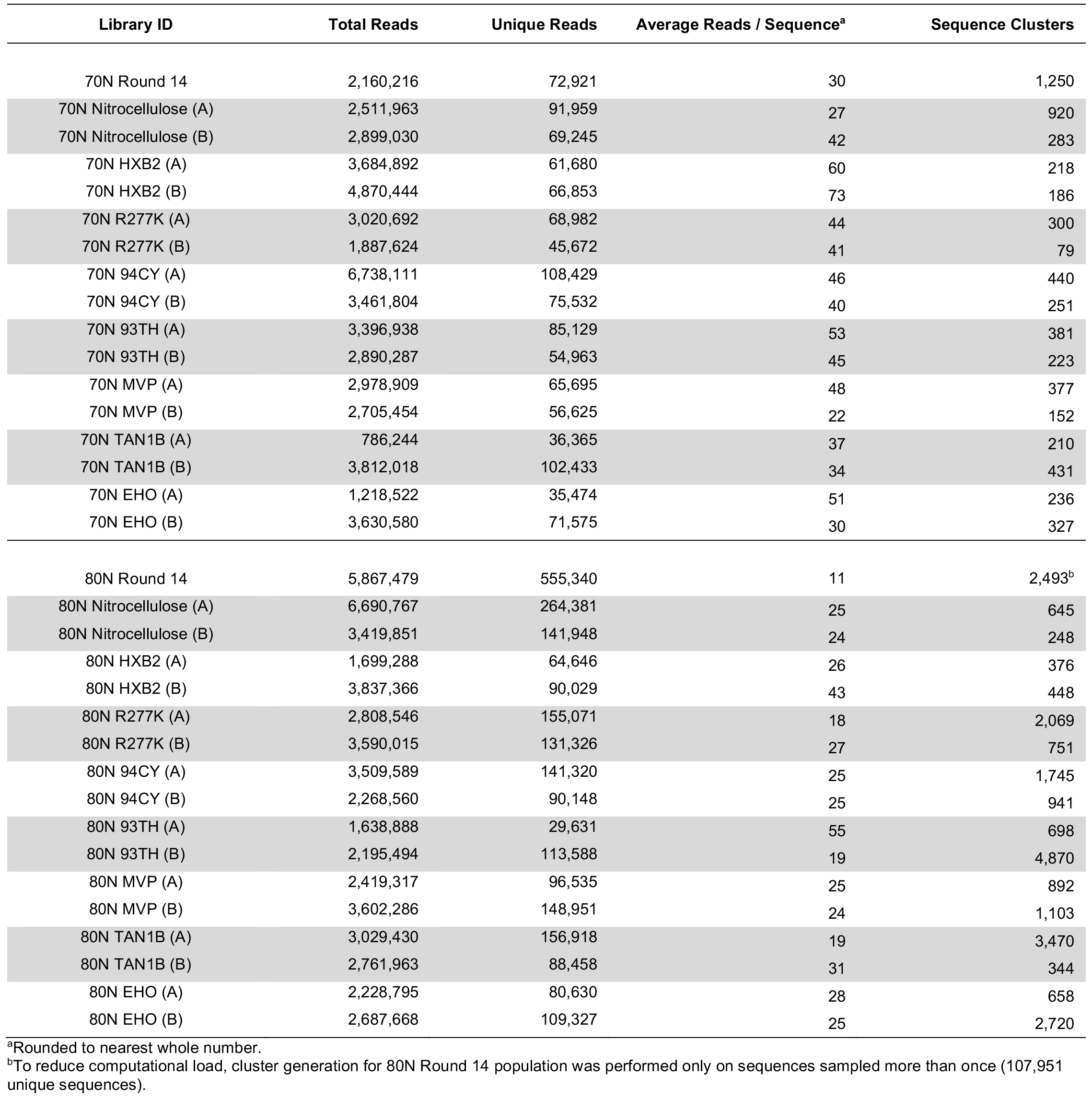
Read counts (total, unique, reads per sequence) and number of unique sequence clusters from high throughput sequencing of Poly-Target populations.

**Figure S3.**
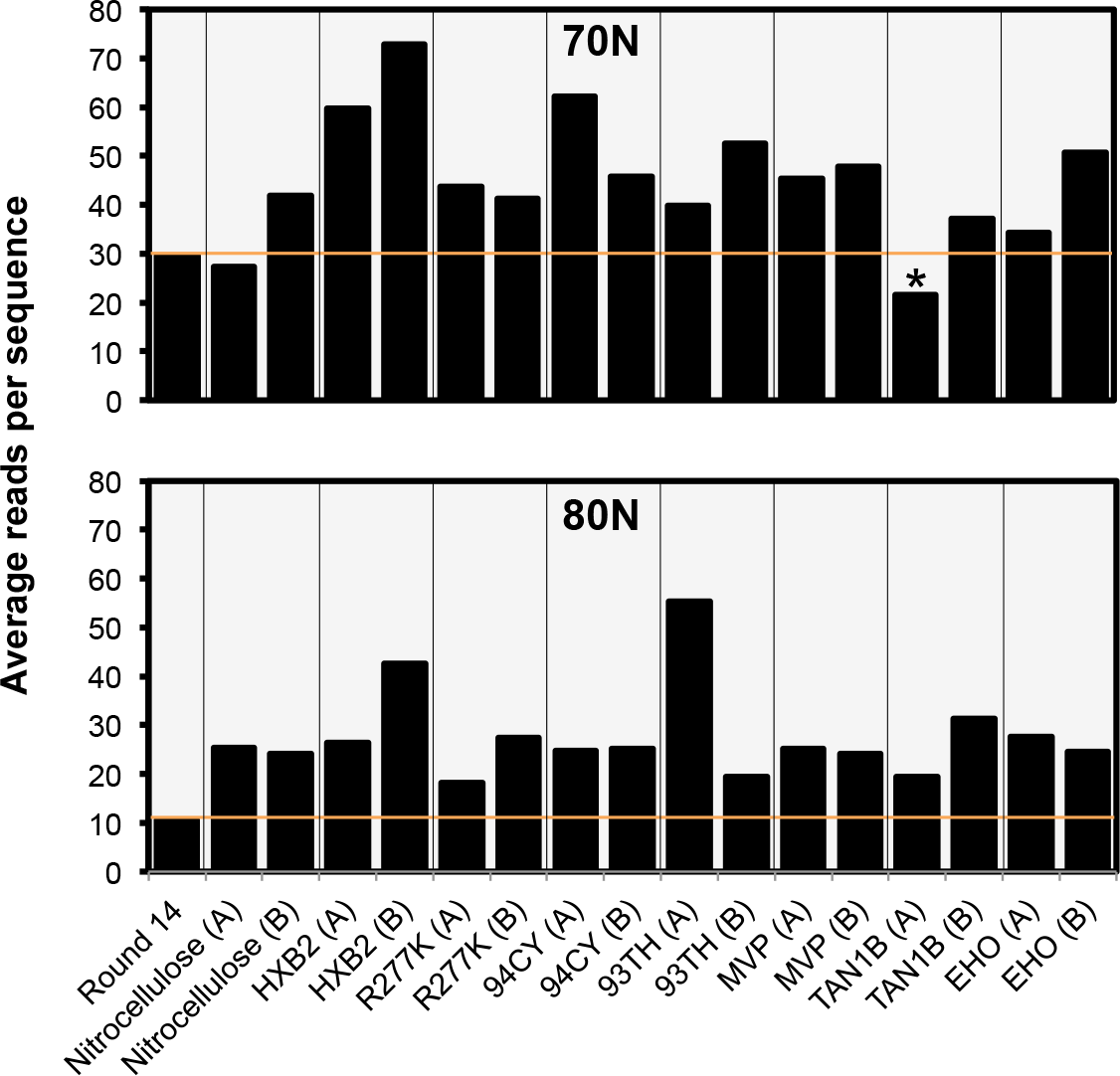
Enrichment of Poly-Target selection trajectories. The average reads per sequence of an aptamer library can be used to monitor the convergence of sequence space of a population when compared to a preceding population. Enrichment was observed in all of the 70N (top) and 80N (bottom) round 17 Poly-Target populations sampled with greater than 1 million reads. A single library, 70N round 17 TAN1B (A), marked with the asterisk had low sampling depth (786,244 reads) and may underrepresent the true diversity of the library relative to the round 14 populations. Orange bars indicate the average reads per sequence of the starting populations for Poly-Target selection.

**Figure S4.**
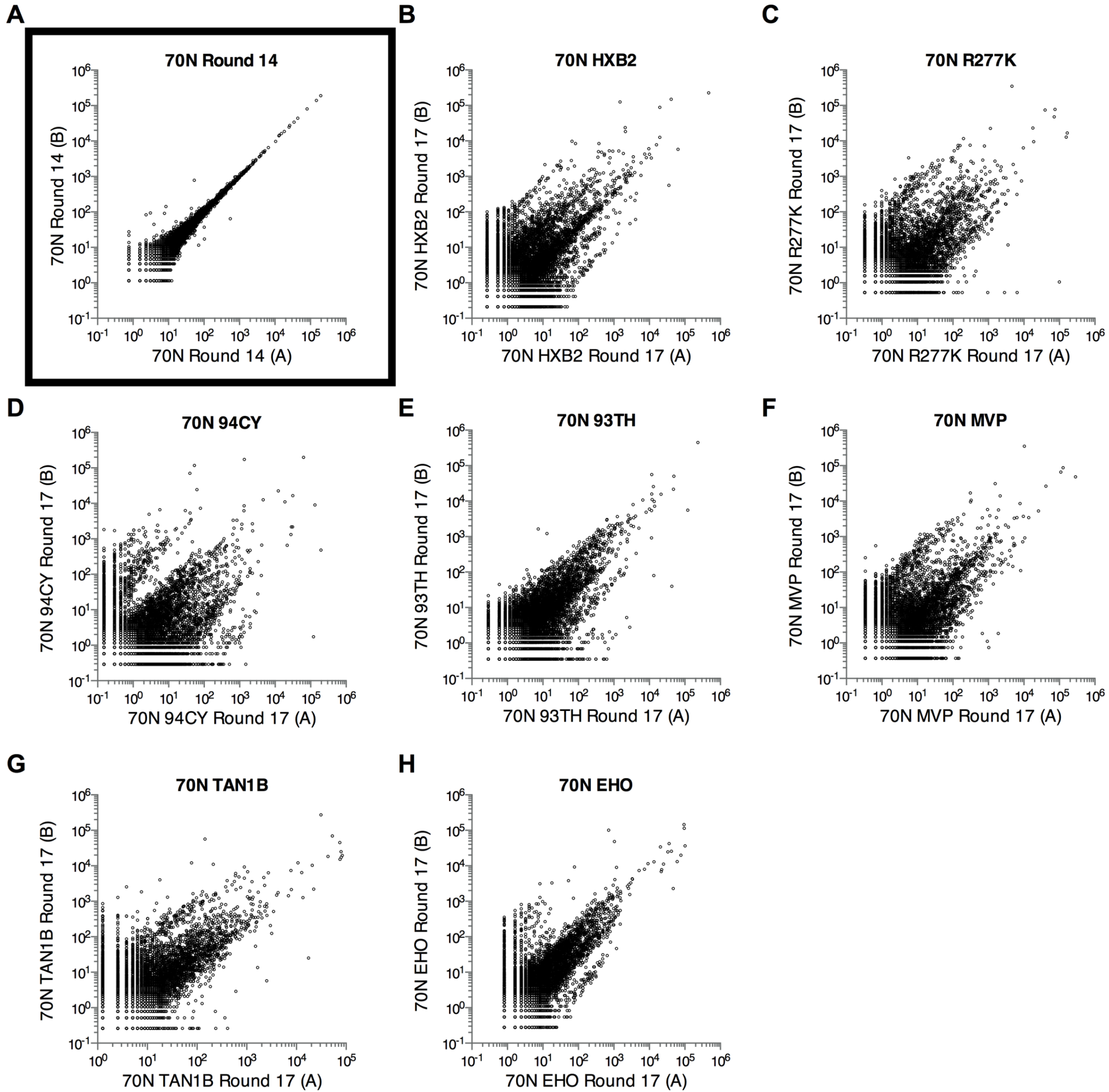
Comparison of 70N replicate trajectories. Comparison of sequence read frequencies in replicate sampling of the 70N round 14 library (**A**), and of replicate selection trajectories after three rounds. (**B**) 70N HXB2 round 17, (**C**) 70N round 17 R277K, (**D**) 70N round 17 94CY, (**E**) 70N round 17 93TH, (**F**) 70N round 17 MVP, (**G**) 70N round 17 TAN1B, (**H**) 70N round 17 EHO. Units of *x-* and *y*-axes are in reads per million and plotted on a logarithmic scale. Sequences sampled only once are not shown.

**Figure S5.**
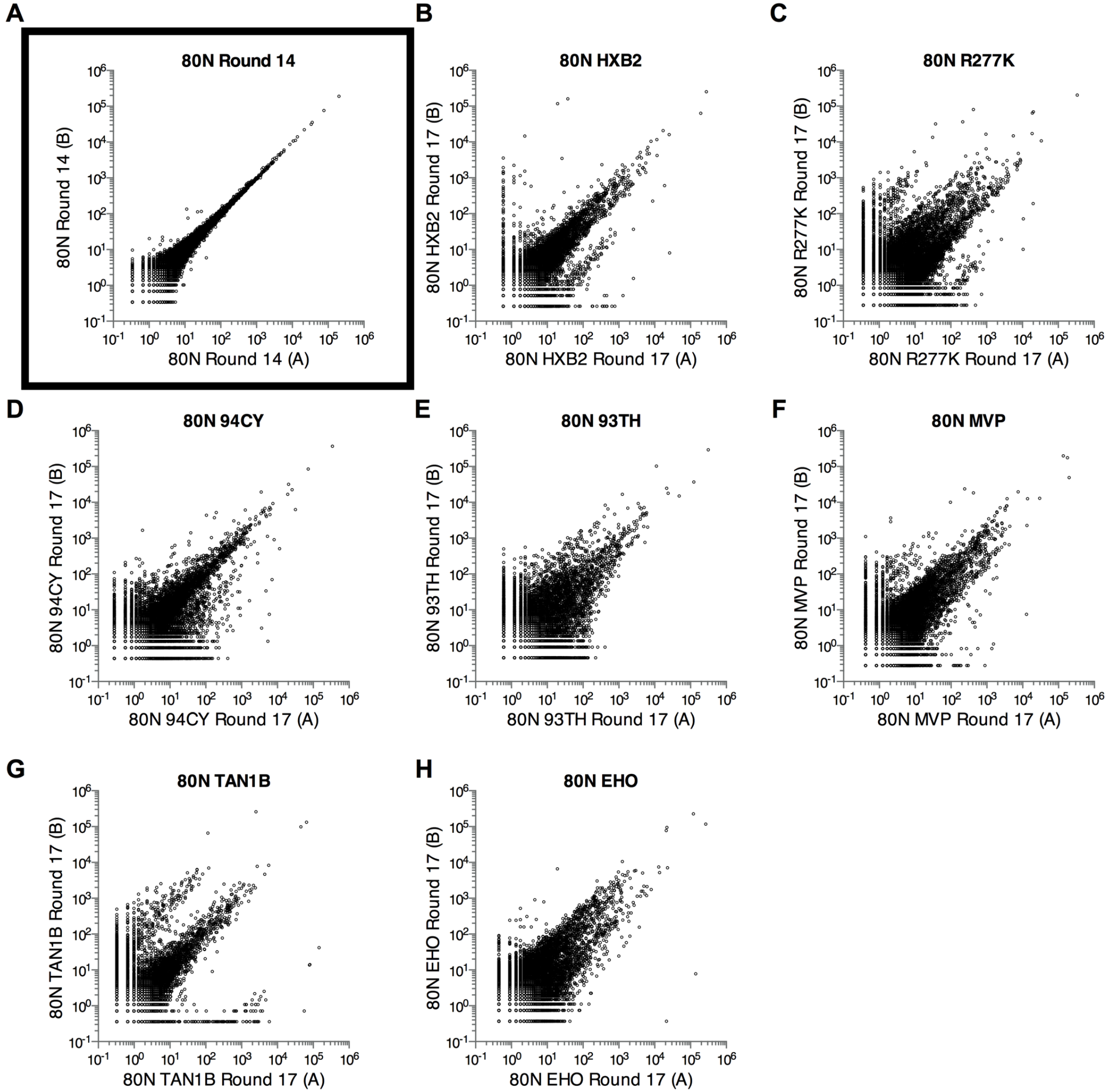
Comparison of 80N replicate trajectories. Comparison of sequence read frequencies in replicate sampling of the 80N round 14 library (**A**), and of replicate selection trajectories after three rounds. (**B**) 80N HXB2 round 17, (**C**) 80N round 17 R277K, (**D**) 80N round 17 94CY, (**E**) 80N round 17 93TH, (**F**) 80N round 17 MVP, (**G**) 80N round 17 TAN1B, (**H**) 80N round 17 EHO. Units of *x-* and *y*-axes are in reads per million and plotted on a logarithmic scale. Sequences sampled only once are not shown.

**Figure S6.**
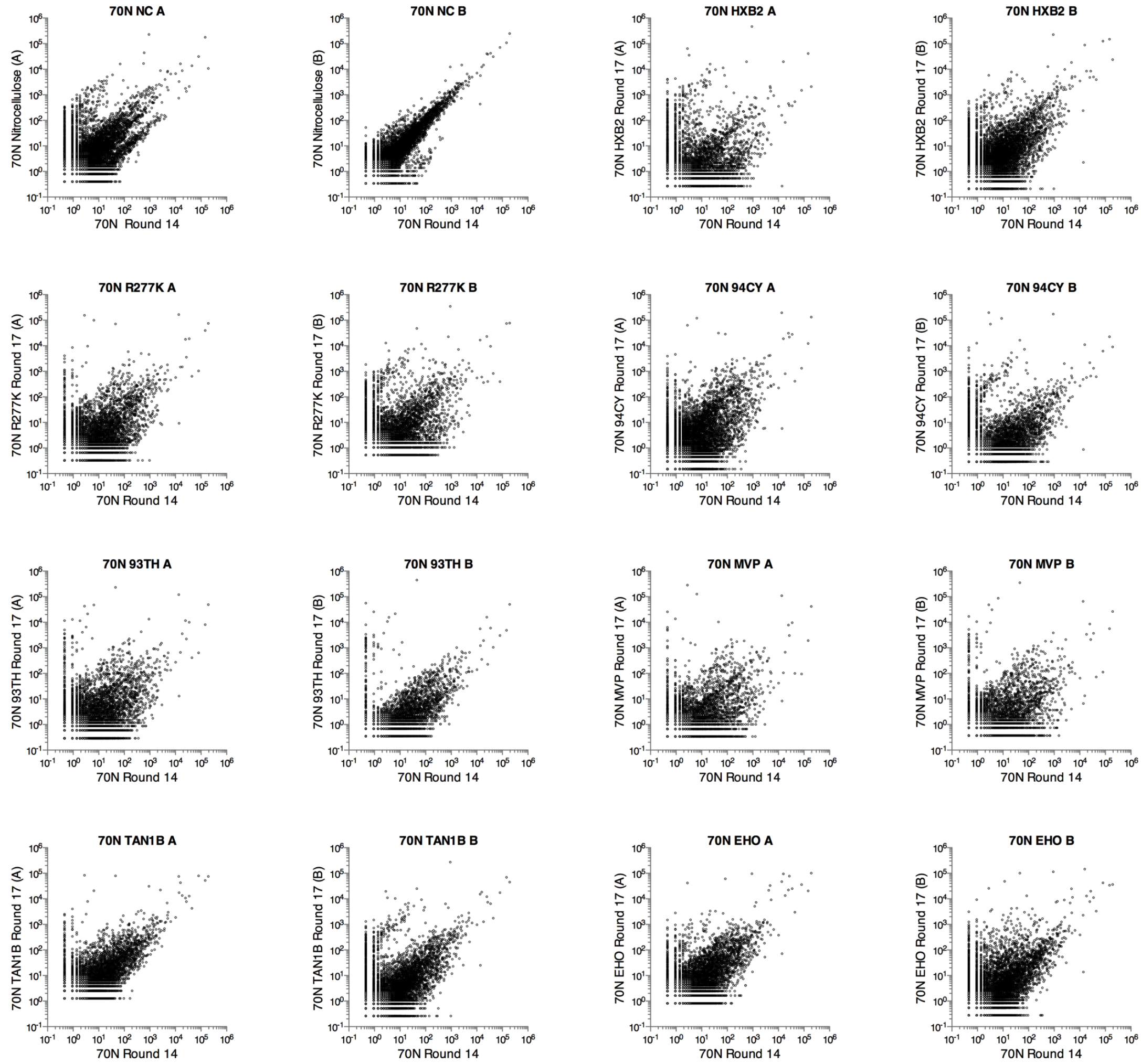
Comparison of 70N round 17 trajectories with the starting round 14 libraries. Comparison of sequence frequencies in 70N round 14 versus nitrocellulose binding population replicates, HXB2 replicates, R277K replicates, 94CY replicates, 93TH replicates, MVP replicates, TAN1B replicates and EHO replicates. Units of *x-* and *y*-axes are in reads per million and plotted on a logarithmic scale. Sequences sampled only once are not shown.

**Figure S7.**
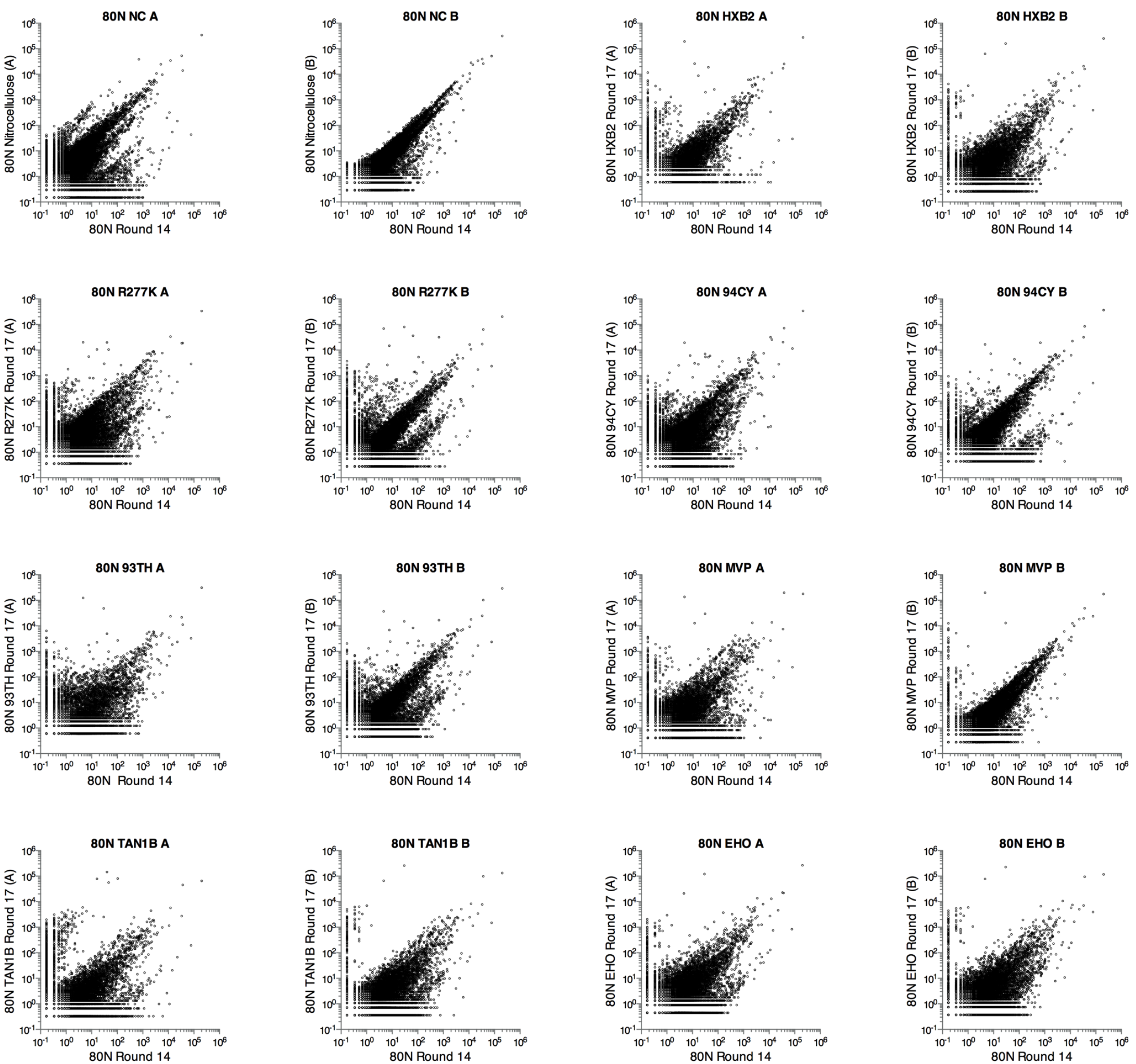
Comparison of 80N round 17 trajectories with the starting round 14 libraries. Comparison of sequence frequencies in 80N round 14 versus nitrocellulose binding population replicates, HXB2 replicates, R277K replicates, 94CY replicates, 93TH replicates, MVP replicates, TAN1B replicates and EHO replicates. Units of *x-* and *y*-axes are in reads per million and plotted on a logarithmic scale. Sequences sampled only once are not shown.

**Figure S8.**
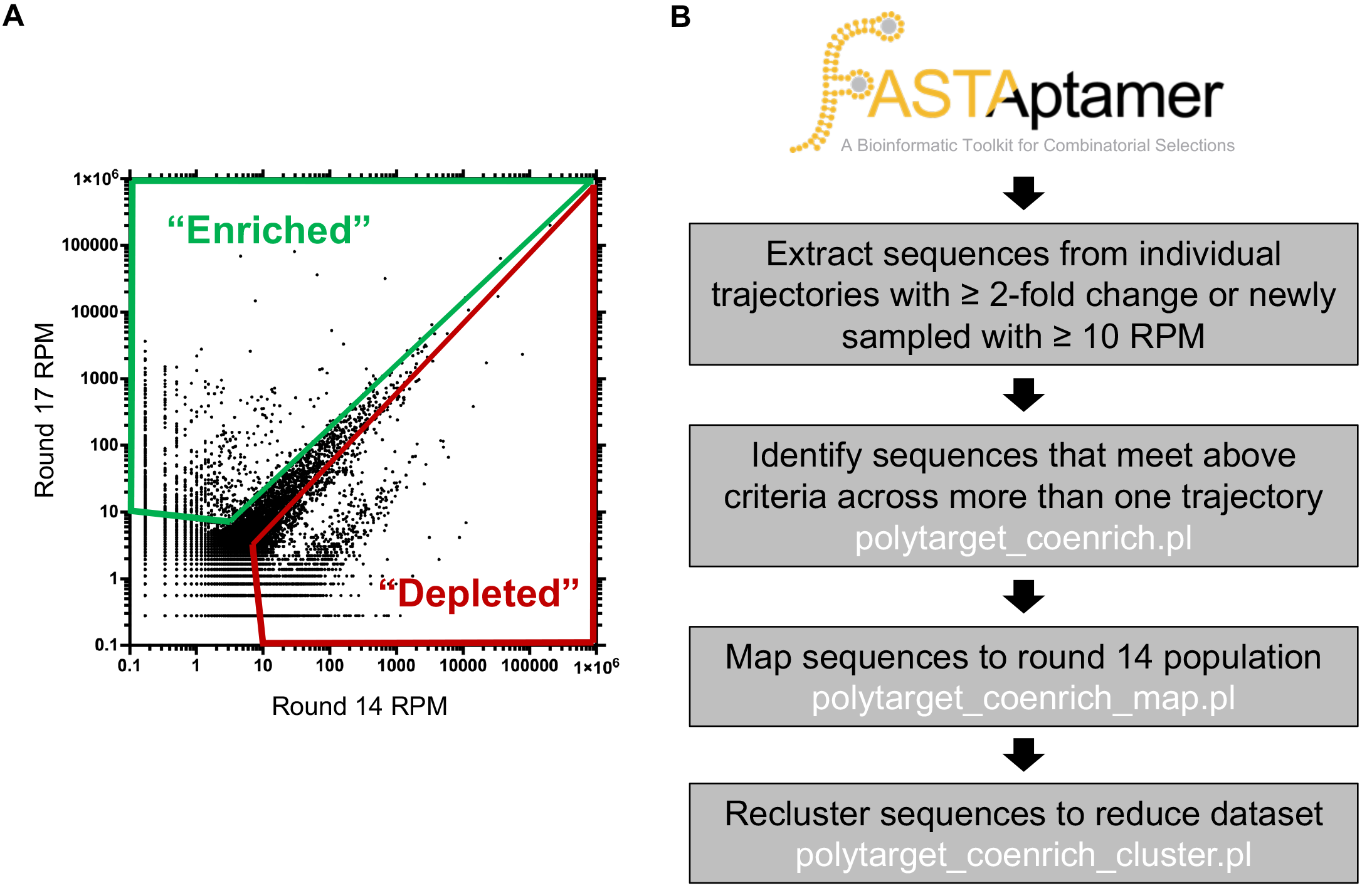
Enrichment and depletion criteria for inclusion into coenrichment analysis. Sequences with at least 10 reads per million (RPM) in aggregate (round 14 RPM + round 17 RPM) were considered in coenrichment analysis. We defined enrichment as ≥ 2-fold increase in sequence RPM (green) and depletion as ≥ 2-fold decrease in sequence RPM from round 14 to round 17 (red). Also included were sequences that were only present at ≥10 RPM in either round 14 (depletion) or round 17 (enrichment) alone, which are not plotted on the graph. (**A**) Shown here is a representative diagram of sequences that would meet the criteria in a single selection trajectory. (**B**) The bioinformatic pipeline for coenrichment analysis began by parsing FASTAptamer-Enrich files for sequences that meet the enrichment or depletion criteria in an individual trajectory (24). Those sequences are then compared across trajectories to identify coenrichers/codepleters, mapped to their round 14 identity (if available), and reclustered to focus characterization on the most represented sequence within a cluster. Perl scripts used for coenrichment analysis are freely available at https://github.com/FASTAptamer/PolyTarget.

**Table S3.**
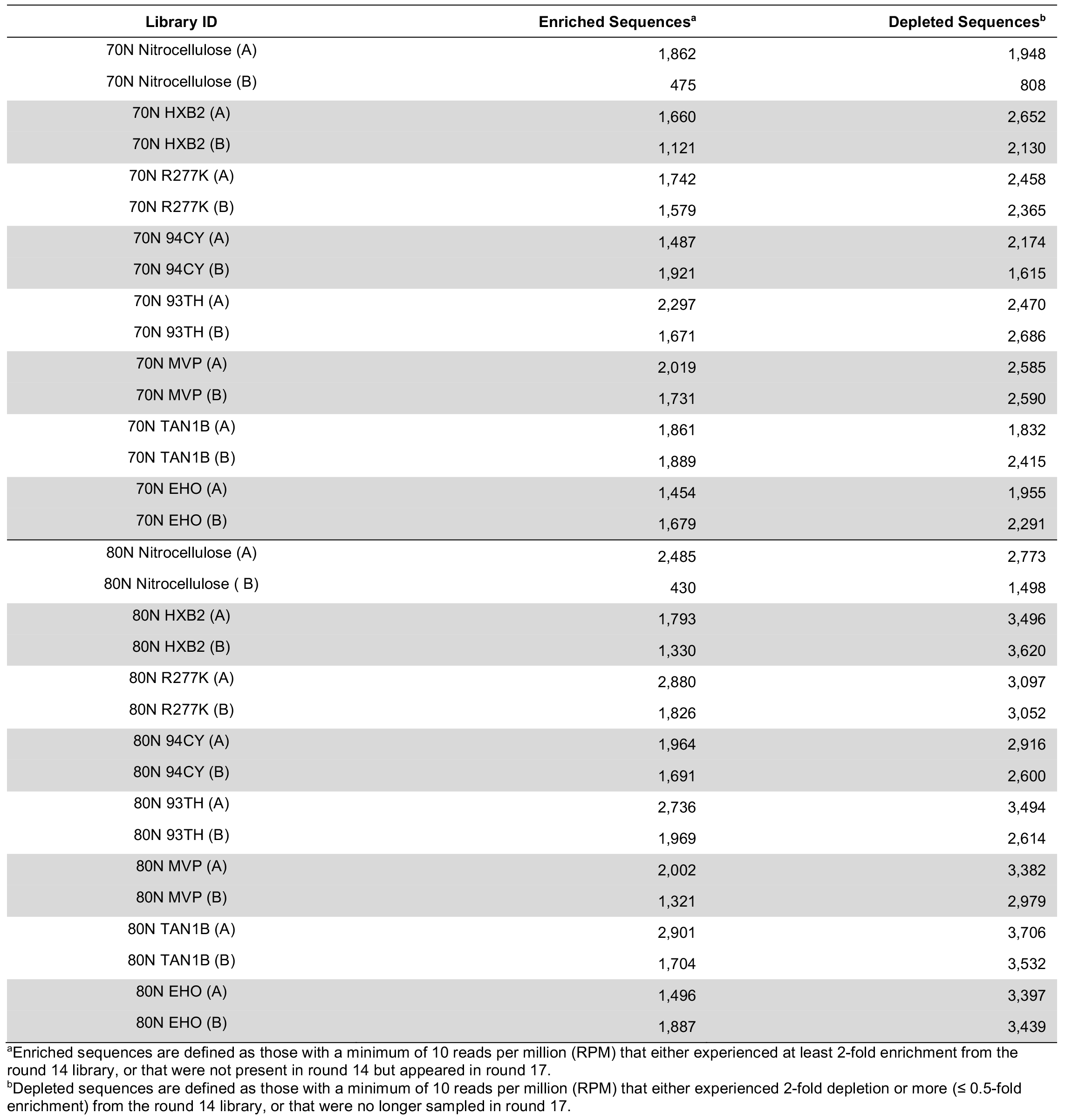
Number of sequences considered enriched or depleted in each Poly-Target selection trajectory.

**Table S4.** Number of sequences that coenriched or codepleted across Poly-Target selection trajectories.

| Library | Coenriched Sequences<sup>a</sup> | Codepleted Sequences<sup>b</sup> |
| --- | --- | --- |
| 70N (A) | 2,129 [3,170] | 1,095 [2,971] |
| 70N (B) | 2,524 [2,778] | 2,222 [2,953] |
| 80N (A) | 2,575 [3,798] | 1,429 [4,145] |
| 80N (B) | 2,823 [2,959] | 2,604 [4,015] |

<sup>a</sup>Coenriched sequences are defined as those that met our criteria for enrichment in at least 2 selection trajectories.
<sup>b</sup>Codepleted sequences are defined as those that met our criteria for depletion in at least 2 selection trajectories.
Values in brackets refer to the number of sequences that coenriched or codepleted in 2 trajectories prior to subtraction of the nitrocellulose population.

**Figure S9.**
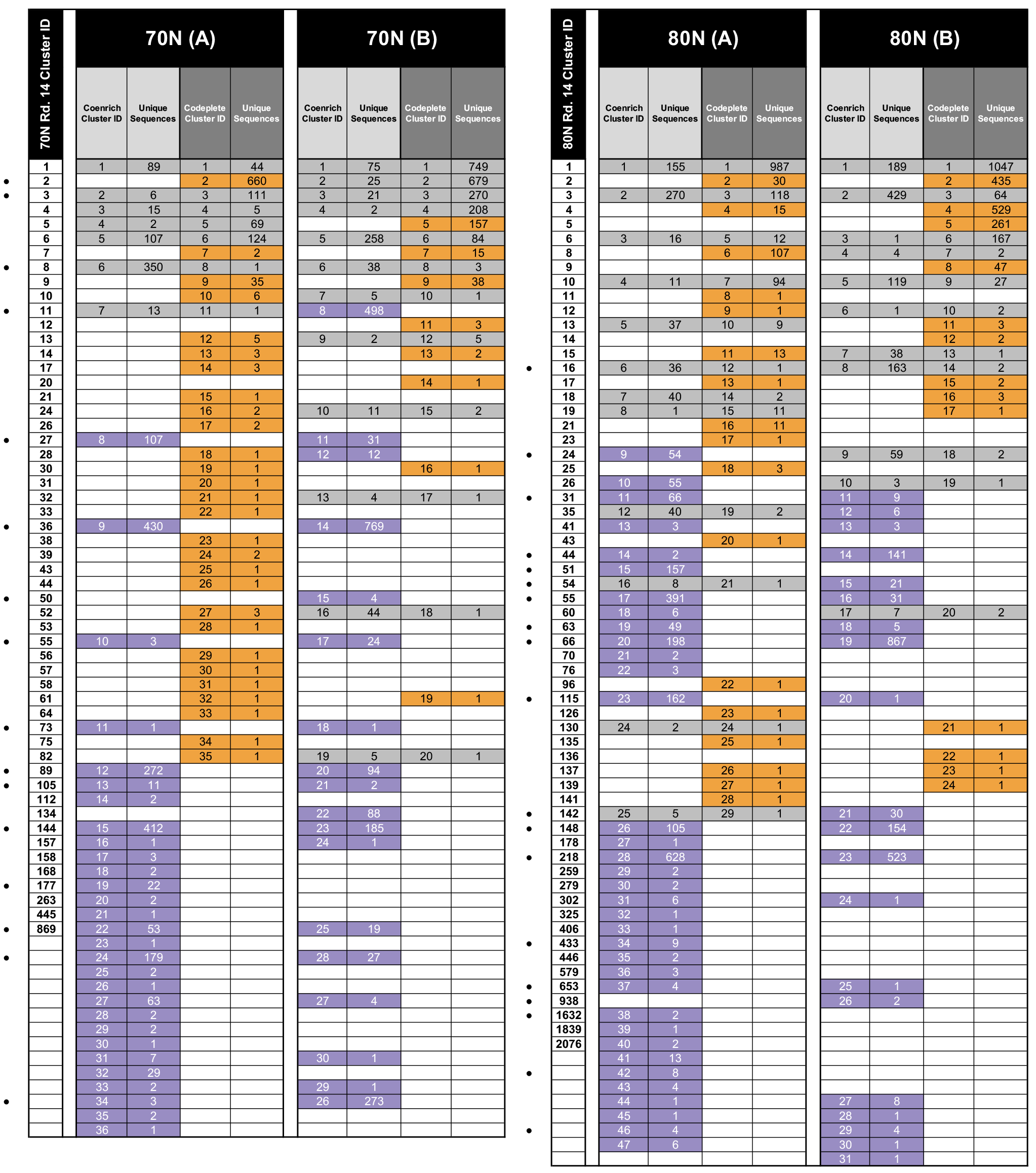
Coenrichment and codepletion analysis of clusters reveal the emergence of rare sequence clusters. Within each replicate trajectory, clusters of related sequences that experienced only depletion are highlighted in gold and those that experienced only enrichment are highlighted in purple. Clusters containing sequences that both enriched and depleted are highlighted in gray. Cluster identities in round 17 populations are traced back to and named after their original cluster identity in the round 14 libraries for comparison within the 70N or 80N subsets. Abundant clusters with large numbers of closely-related sequences include individual cluster members that score as co-enriching and others that score as co-depleting, such that the cluster as a whole is subject to both depletion and enrichment. Clusters that were rare in round 14 almost exclusively experienced enrichment. Dots (•) to the left of cluster designations indicate clusters that were chosen for further analysis.

**Figure S10.**
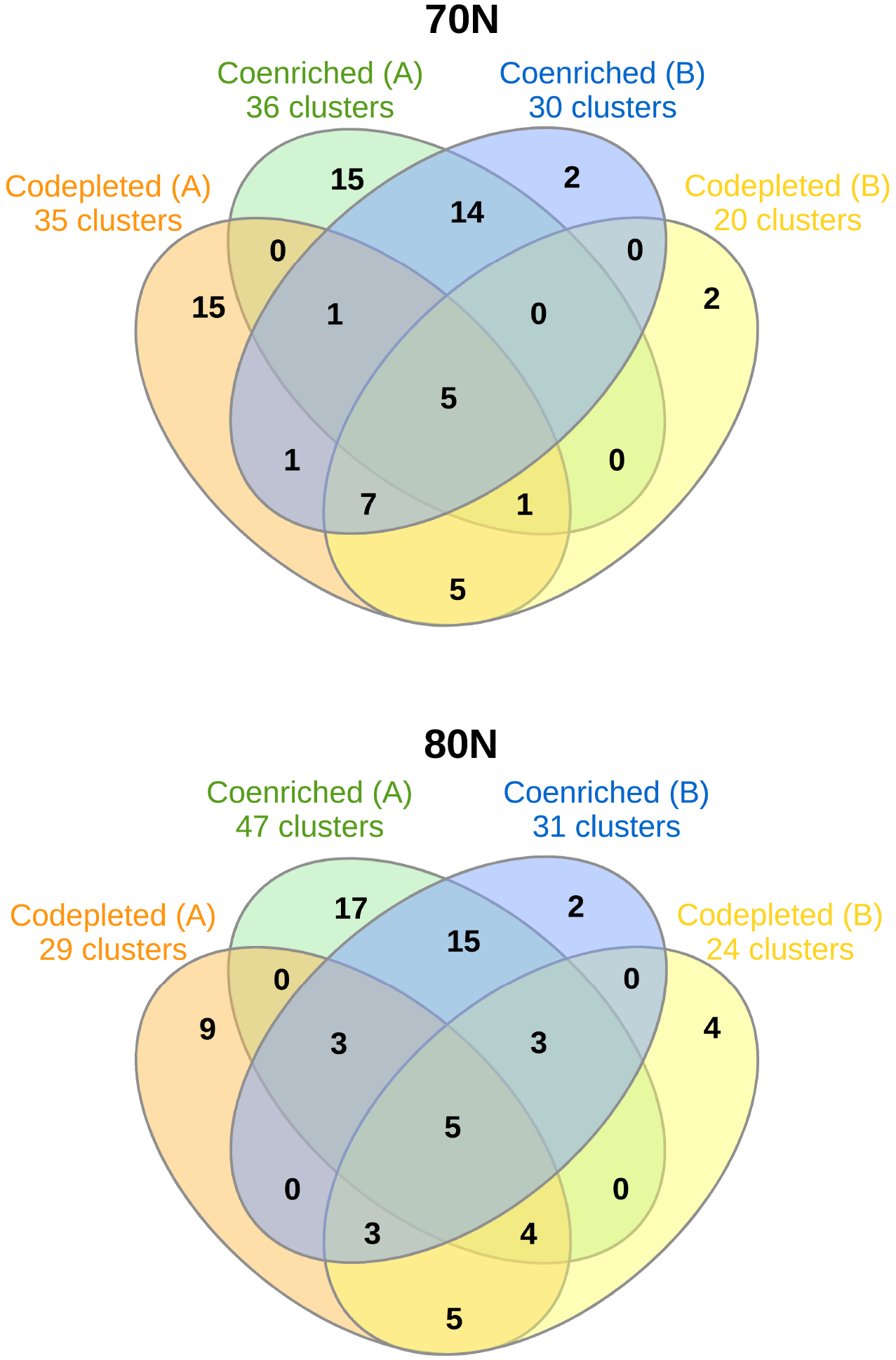
Set diagram of coenriched and codepleted clusters from the 70N and 80N selections. Selection of candidate aptamers was informed by the identification of clusters that experienced coenrichment and/or codepletion from the 70N (top) and 80N (bottom) libraries. Of particular interest were the clusters that experienced only coenrichment in either or both of the replicates.

**Table S5.**
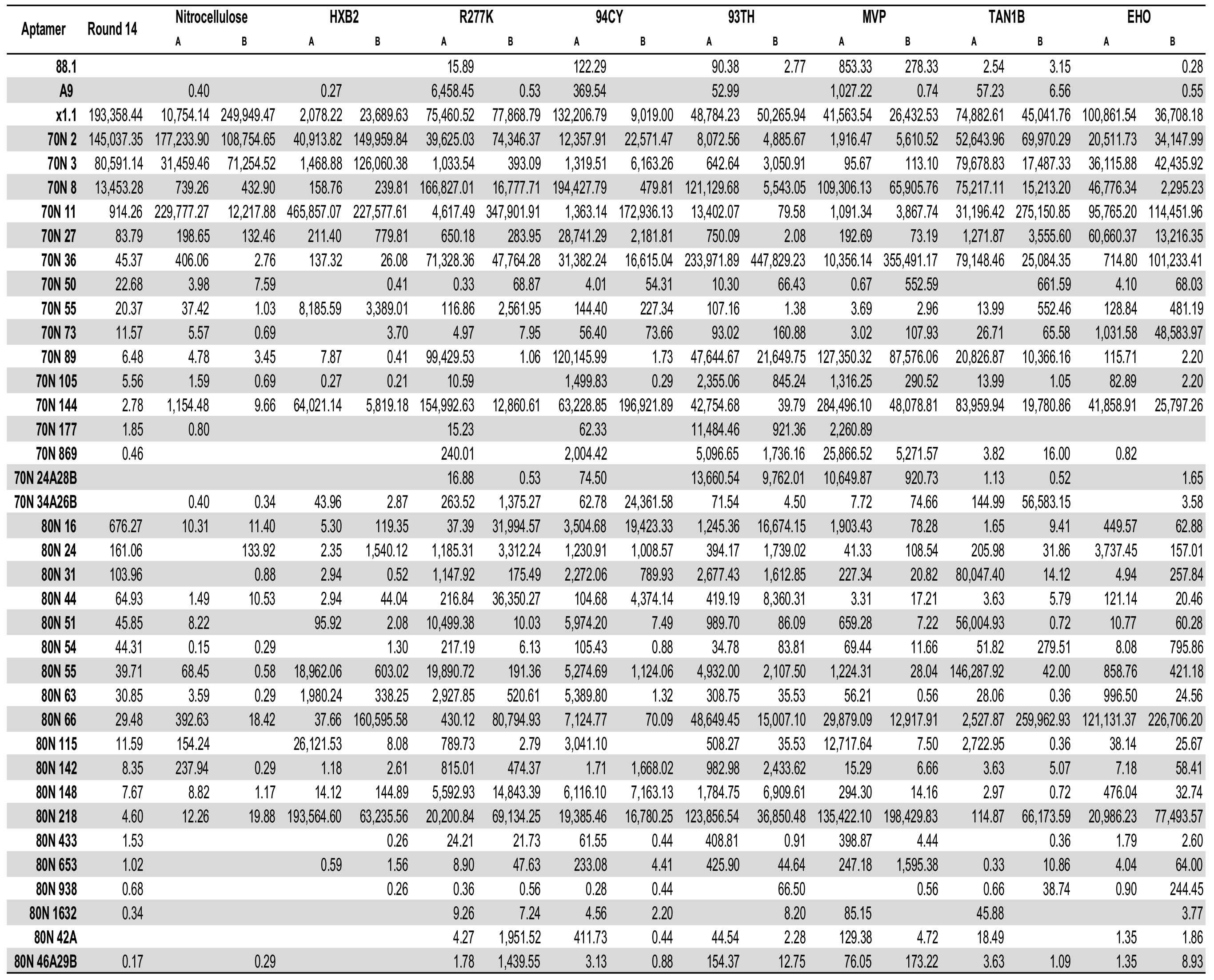
Reads per million of candidate aptamers identified through Poly-Target selection and coenrichment/cordepletion analysis.

**Figure S11.**
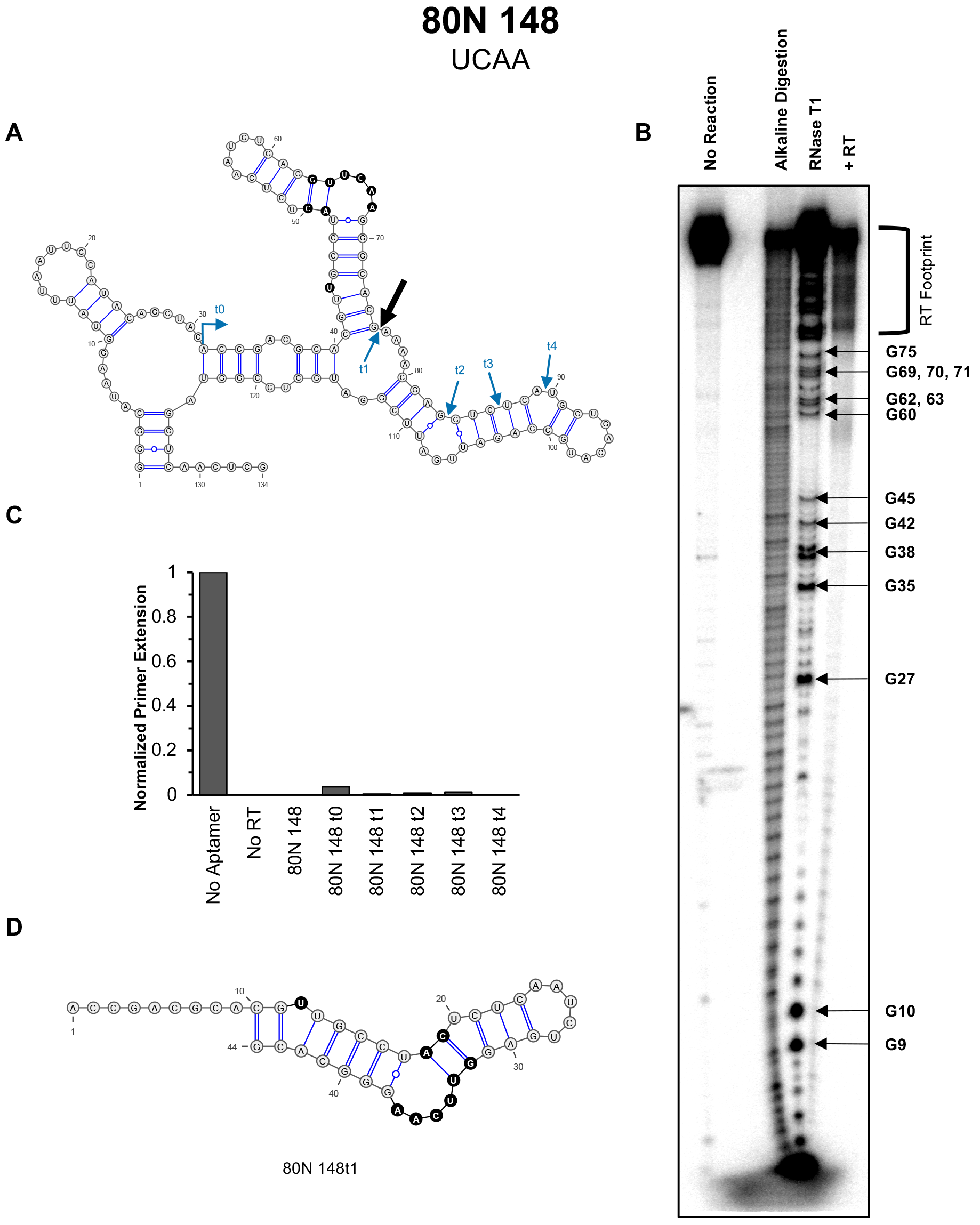
Minimization and motif characterization of 80N 148. (**A**) The predicted secondary structure of the aptamer with arrows depicting the location of the 3’ boundary using RNase T1 digestion (large black arrow), and 3’ end truncations that either remain functional (blue) or abrogate inhibition (red). (**B**) RT:aptamer 3’ boundary. (**C**) Inhibition of HXB2 RT’s DNA-dependent DNA polymerase activity by aptamer truncations. (**D**) The minimally viable 3’ end truncation of the aptamer, 80N 148t1.

**Figure S12.**
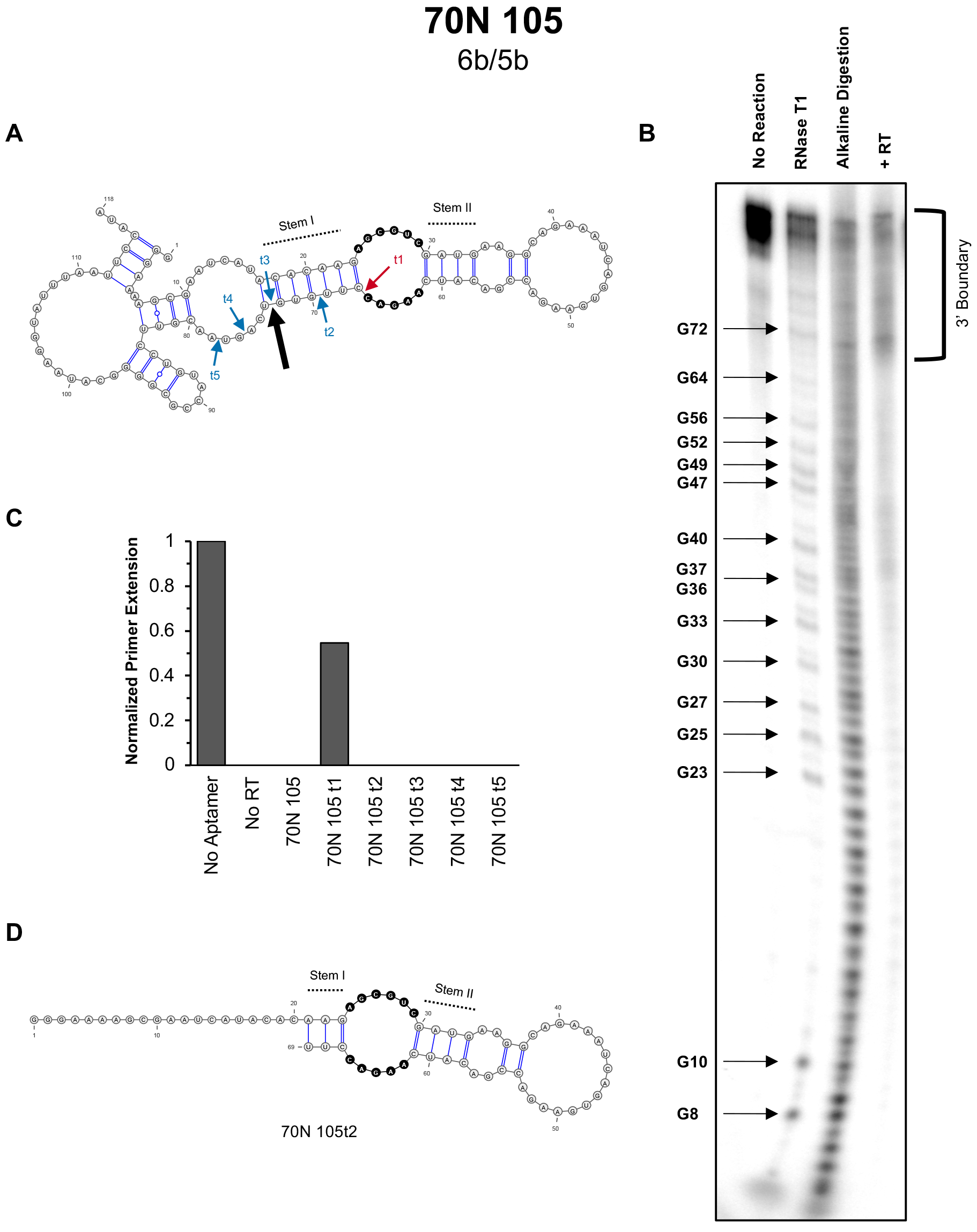
Minimization and motif characterization of 70N 105. (**A**) The predicted secondary structure of the aptamer with arrows depicting the location of the 3’ boundary using RNase T1 digestion (large black arrow), and 3’ end truncations that either remain functional (blue) or abrogate inhibition (red). (**B**) RT:aptamer 3’ boundary. (**C**) Inhibition of HXB2 RT’s DNA-dependent DNA polymerase activity by aptamer truncations. (**D**) The minimally viable 3’ end truncation of the aptamer, 70N 105t2.

**Figure S13.**
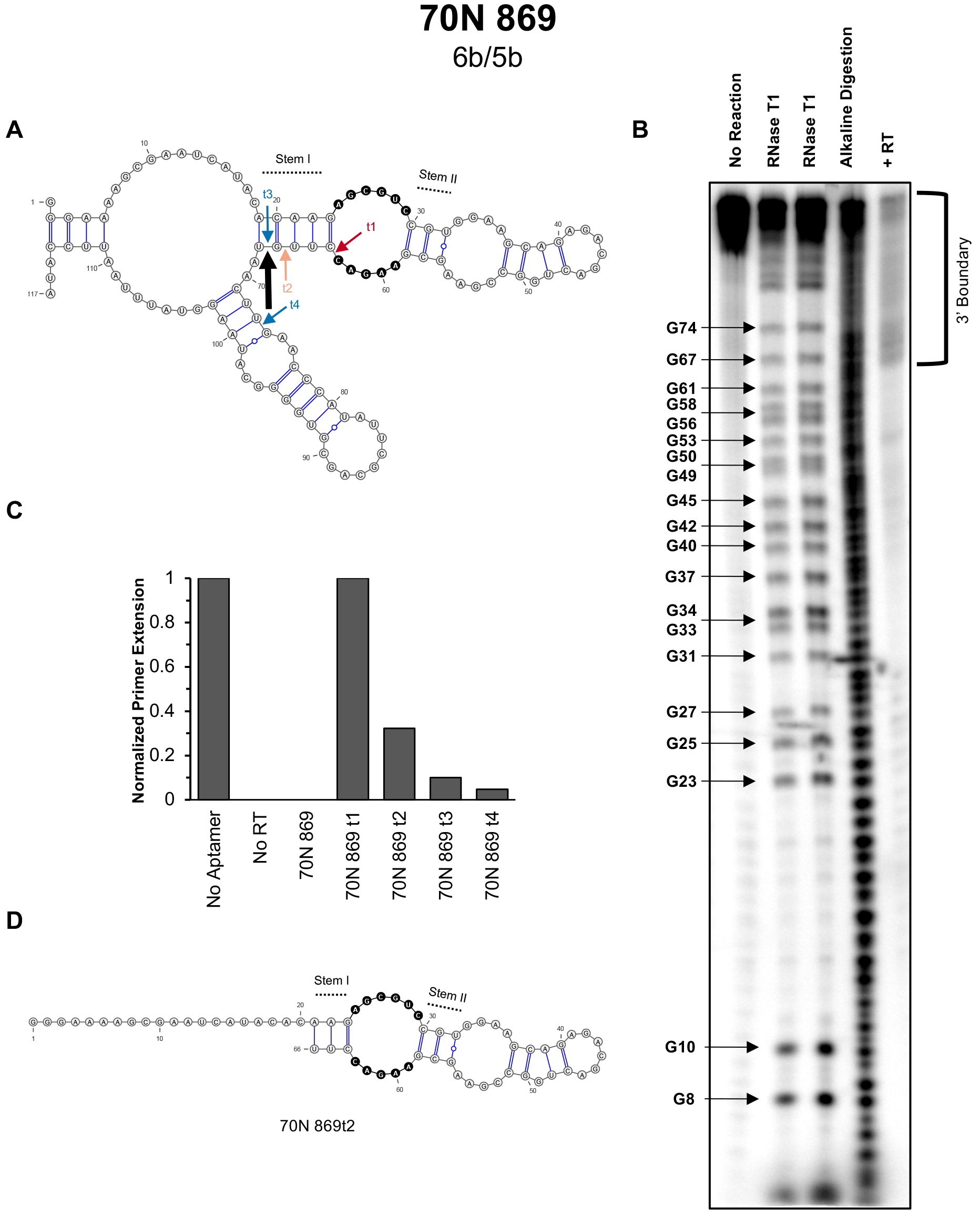
Minimization and motif characterization of 70N 869. (**A**) The predicted secondary structure of the aptamer with arrows depicting the location of the 3’ boundary using RNase T1 digestion (large black arrow), and 3’ end truncations that either remain functional (blue) or abrogate inhibition (red). (**B**) RT:aptamer 3’ boundary. (**C**) Inhibition of HXB2 RT’s DNA-dependent DNA polymerase activity by aptamer truncations. (**D**) The minimally viable 3’ end truncation of the aptamer, 70N 869t2.

**Figure S14.**
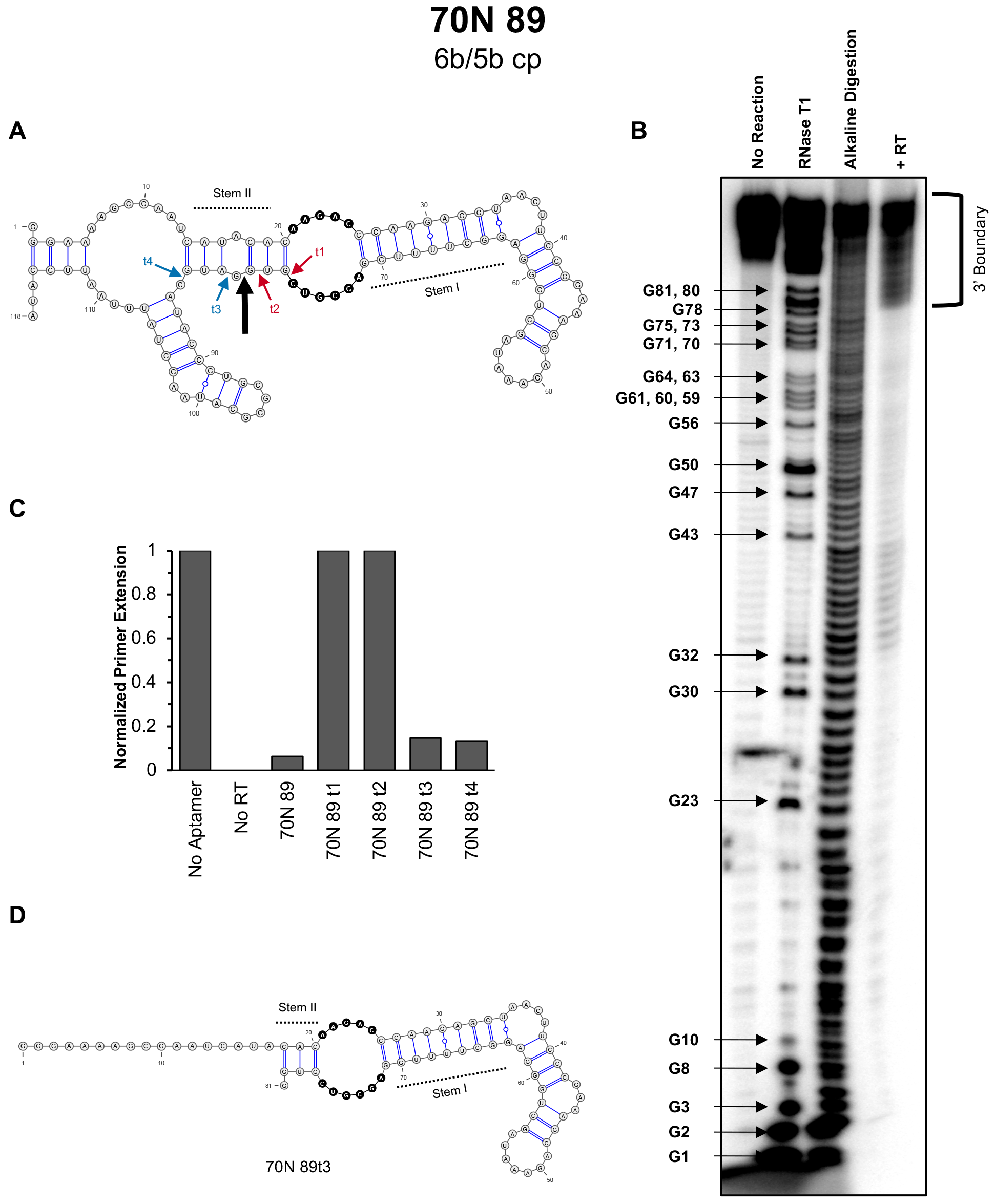
Minimization and motif characterization of 70N 89. (**A**) The predicted secondary structure of the aptamer with arrows depicting the location of the 3’ boundary using RNase T1 digestion (large black arrow), and 3’ end truncations that either remain functional (blue) or abrogate inhibition (red). (**B**) RT:aptamer 3’ boundary. (**C**) Inhibition of HXB2 RT’s DNA-dependent DNA polymerase activity by aptamer truncations. (**D**) The minimally viable 3’ end truncation of the aptamer, 70N 89t3.

**Figure S15.**
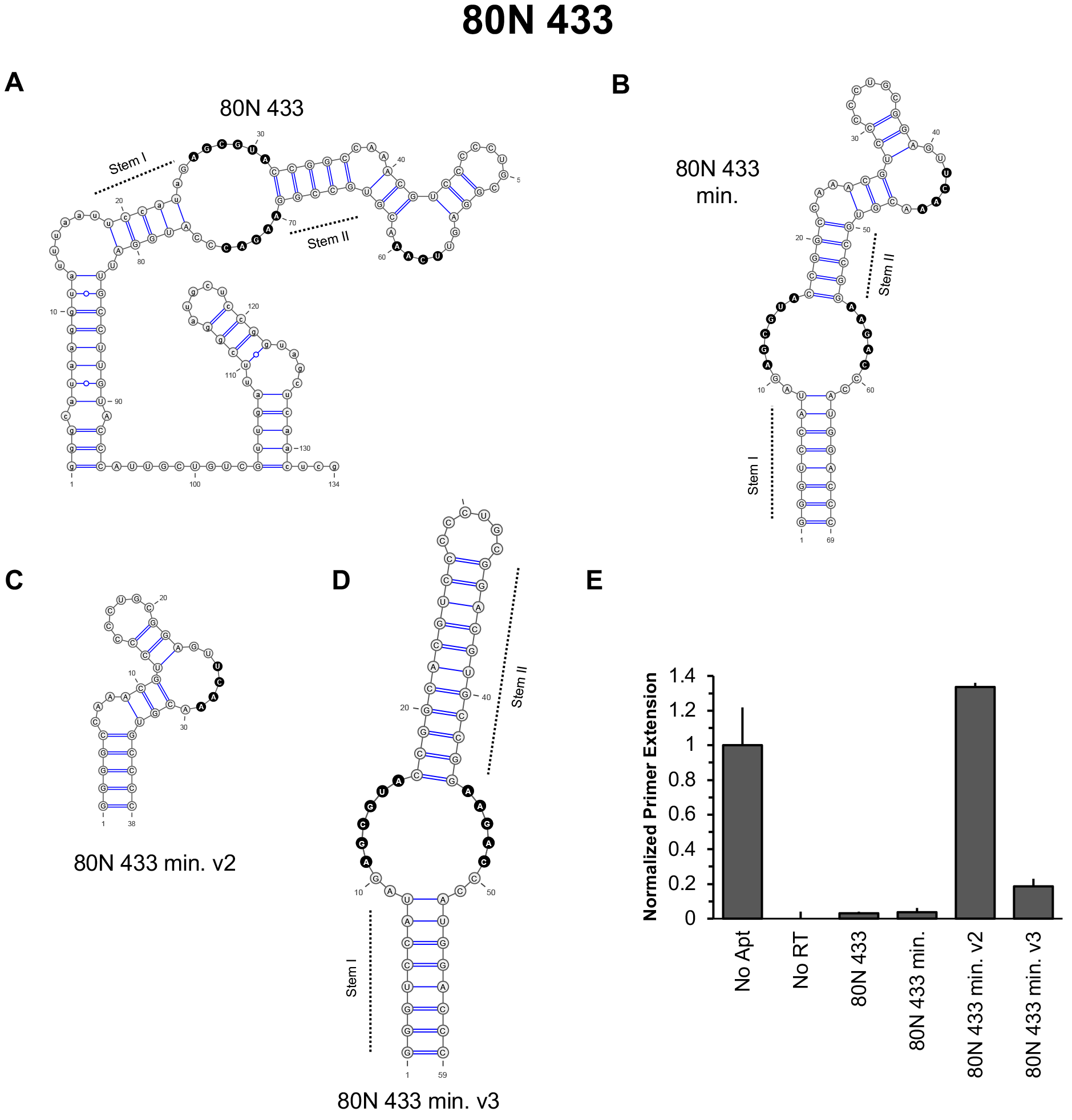
Minimization and motif characterization of 80N 433. (**A**) The predicted secondary structure of full-length aptamer 80N 433 potentially contains both (6/5)AL and UCAA bulge motifs. Note that Stem I can be extended by two more base pairs if A24 and C76 are tolerated in the penultimate closing position to generate asymmetric loops of the expected 6/5 size. (**B**) 80N 433 min., a minimized variant of 80N 433, contains a 5’ GGG to increase transcriptional yield using T7 RNA polymerase and a corresponding 3’ CCC to base pair. (**C**) 80N 433 min. v2 is a variant of the minimal aptamer in which the (6/5)AL motif has been removed, leaving only the bulges. (**D**) 80N 433 min. v3 is a variant of the minimal aptamer in which the predicted CAA and UCAA bulges have been removed, leaving only the (6/5)AL motif. (**E**) Primer extension assays against HXB2 RT. The minimal aptamer, 80N 433 min., shows similar inhibition to the full-length aptamer. Deletion of the (6/5)AL motif in 80N 433 min. v2 abrogates the inhibitory function of the aptamer. Removal of only the bulges (including the UCAA-containing bulge) reduces the primer extension ability of HXB2 RT but remains less inhibitory than the variants containing both the bulges and the (6/5)AL motif.

**Figure S16.**
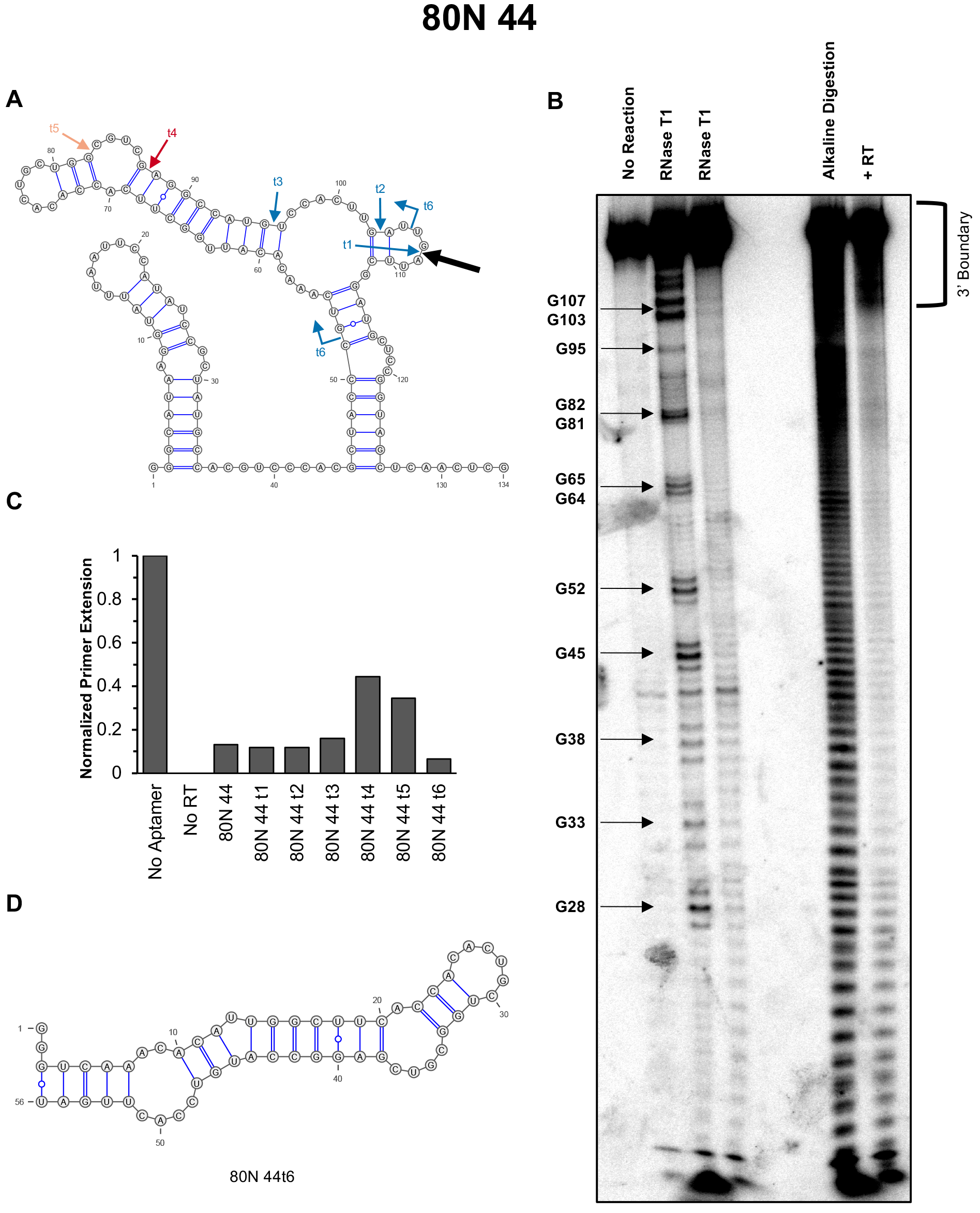
Minimization and motif characterization of 80N 44. (**A**) The predicted secondary structure of the aptamer with arrows depicting the location of the RT footprint using RNase T1 digestion (large black arrow), and 3’ end truncations that either remain functional (blue) or lower inhibition (orange/red). Truncation 6 (t6) is truncated from both 5’ and 3’ ends. (**B**) RT:aptamer 3’ boundary. (**C**) Inhibition of HXB2 RT’s DNA-dependent DNA polymerase activity by aptamer truncations. (**D**) The minimally viable 3’ end truncation of the aptamer, 80N 44t6.

**Figure S17.**
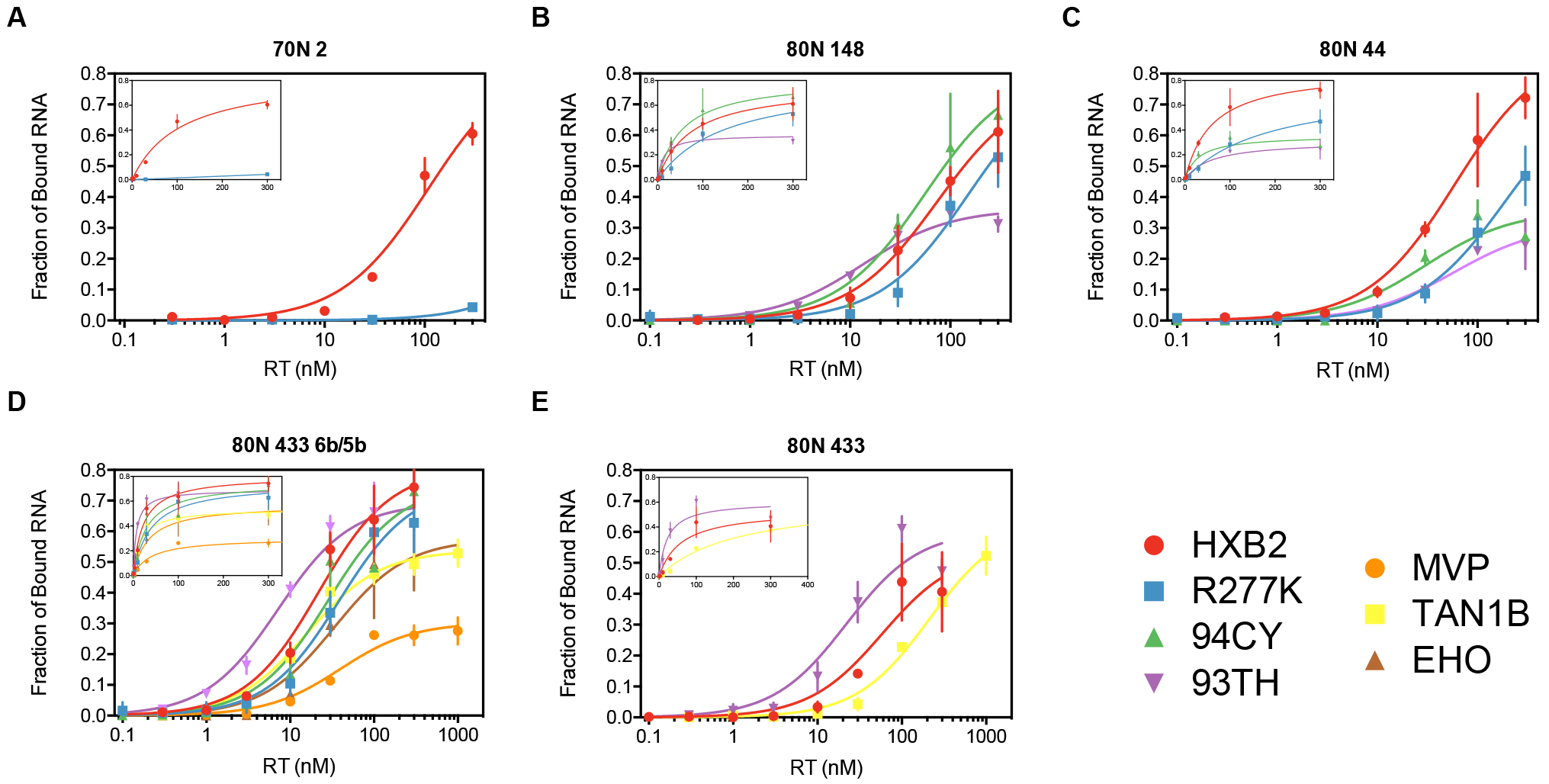
Binding curves for candidate aptamers against various RTs. Trace amounts of 5’ radiolabeled aptamers were incubated with varying concentrations of the indicated RTs. Retention of radiolabel after nitrocellulose filtration was measured and data were fit to a one site binding curve using Prism GraphPad 6.2. Data shown are from triplicate experiments with error bars to indicate standard deviation. Insets show the same data plotted with a linear *x*-axis.

**Table S6.**
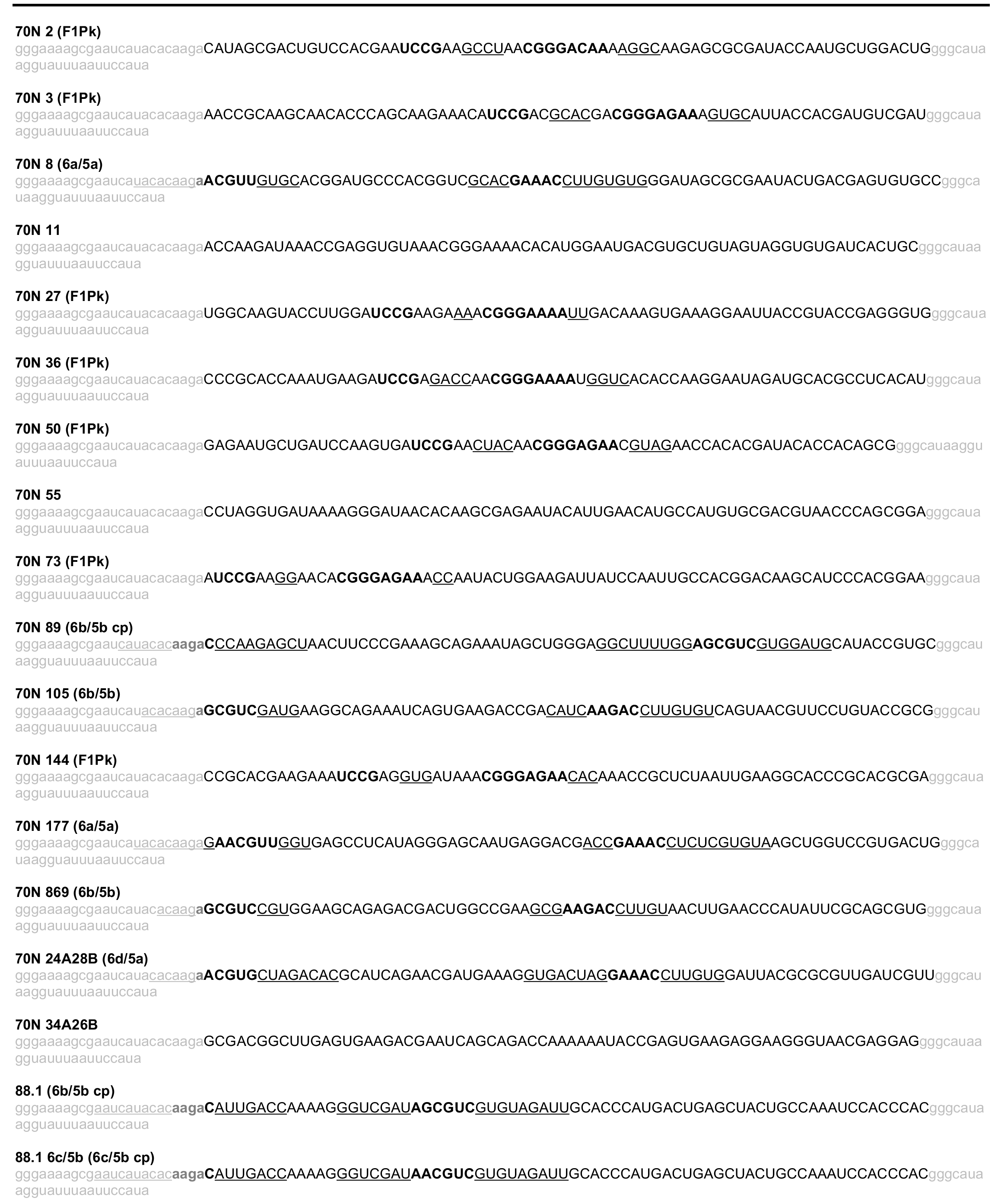
Aptamer sequences used in this study^a^

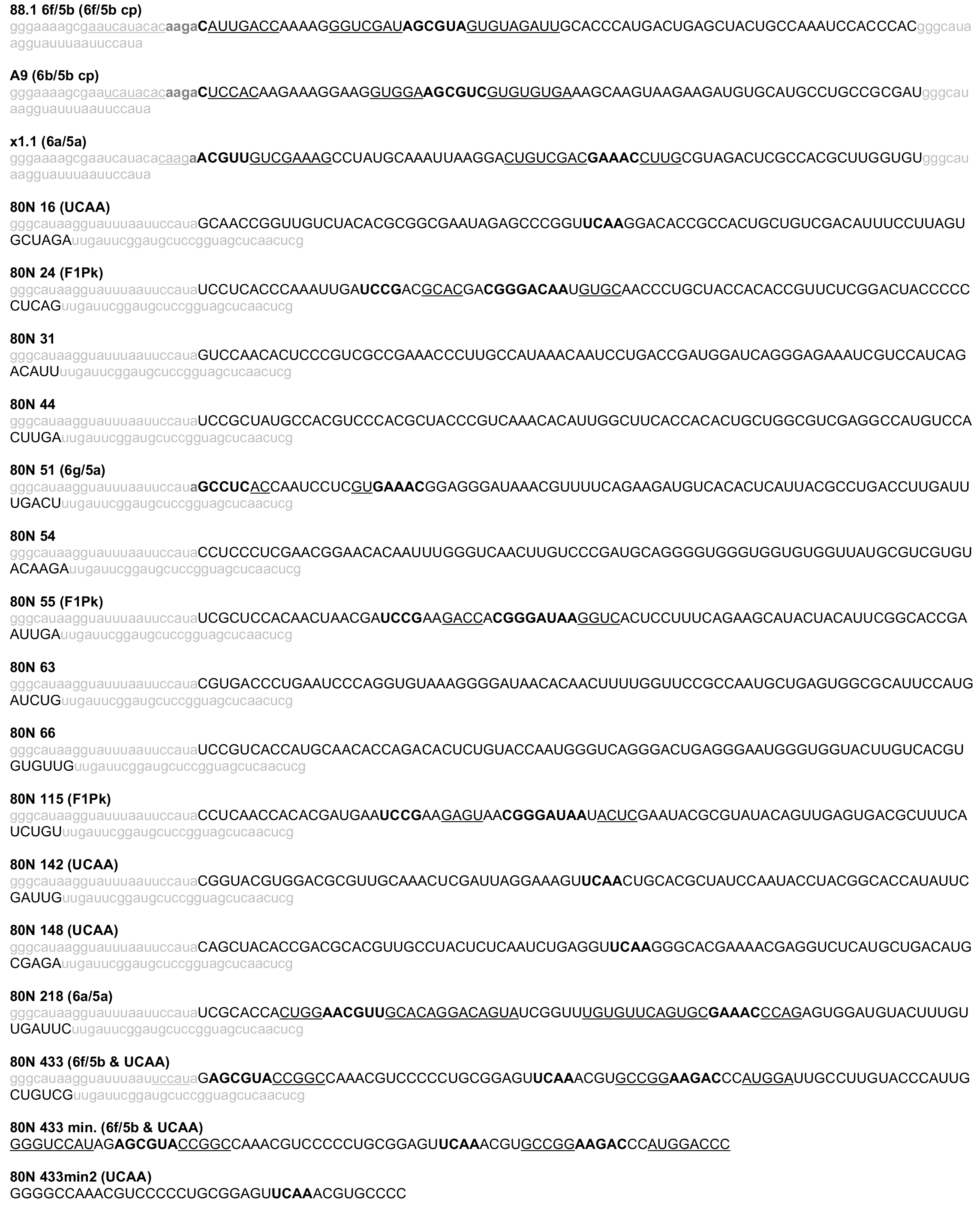

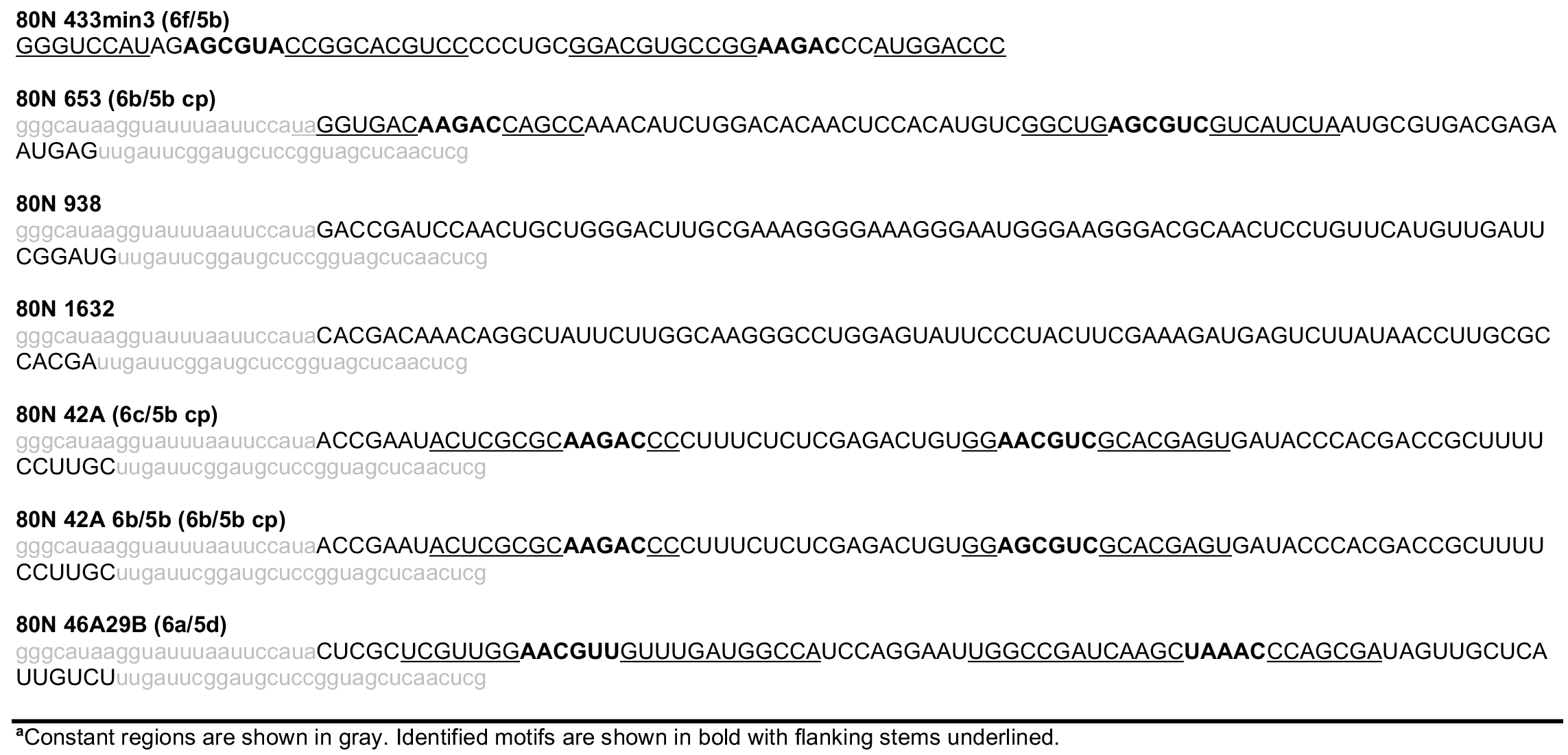

